# Endopiriform neurons projecting to ventral CA1 are a critical node for recognition memory

**DOI:** 10.1101/2024.03.17.585387

**Authors:** Naoki Yamawaki, Hande Login, Solbjørg Østergaard Feld-Jakobsen, Bernadett Mercedesz Molnar, Mads Zippor Kirkegaard, Maria Moltesen, Aleksandra Okrasa, Jelena Radulovic, Asami Tanimura

**Affiliations:** Department of Biomedicine, Aarhus University, Aarhus C, Denmark; PROMEMO, The Center for Proteins in Memory, Aarhus University, Aarhus C, Denmark; DANDRITE, The Danish Research Institute of Translational Neuroscience, Aarhus University, Aarhus C, Denmark; Dominick P. Purpura Department of Neuroscience, Albert Einstein College of Medicine, New York, USA; Department of Psychiatry and Behavioral Sciences, Albert Einstein College of Medicine, New York, USA

## Abstract

The claustrum complex is viewed as fundamental for higher order cognition; however, the circuit organization and function of its neuroanatomical subregions are not well understood. We demonstrated that some of the key roles of the CLA complex can be attributed to the connectivity and function of a small group of neurons in its ventral subregion, the endopiriform (EN). We identified a subpopulation of EN neurons by their projection to the ventral CA1 (EN^vCA1-proj.^ neurons), embedded in recurrent circuits with other EN neurons and the piriform cortex. Although the EN^vCA1-proj.^ neuron activity was biased toward novelty across stimulus categories, their chemogenetic inhibition selectively disrupted the memory-guided but not innate responses of mice to novelty. Based on our functional connectivity analysis, we suggest that EN^vCA1-proj.^ neurons serve as an essential node for recognition memory through recurrent circuits mediating sustained attention to novelty, and through feed forward inhibition of distal vCA1 neurons shifting memory-guided behavior from familiarity to novelty.

## Introduction

The claustrum (CLA) complex, an evolutionarily conserved brain region found across mammalian species, reptiles, and birds, is hypothesized to serve as a node for establishing higher order cognitive functions by coordinating neuronal activity on a global scale (*1–3*). This view is based on its extensive connections with many cortical and subcortical areas (*4, 5*). Some of these functions include sensory perception and attention, which may affect memory processing, including working memory, associative memory, and recognition memory (*6–8*). Consistent with this view, abnormalities of the CLA complex is found in major cognitive disorders, including Alzheimer’s disease, schizophrenia, and attention-deficit/hyperactivity disorders (*9–11*). However, microcircuitry and functional segregation of the individual constituents of the CLA complex have remained unclear.

The cytoarchitecture and genetic expression broadly divide the CLA complex into dorsal and ventral parts, with the dorsal part typically referred to as “CLA” and ventral part having its own nomenclature: “endopiriform (EN)” in rodents (*12, 13*). Current evidence indicates that the CLA forms reciprocal connections with dorsal cortices containing sensory, motor and association areas generating diverse physiological effects depending on the cortical targets and their activity patterns (*4, 5*). In contrast, EN primarily provides inputs to the piriform cortex and limbic systems (*14, 15*), suggesting its distinct circuit organization and role in bridging olfactory information processed by the piriform cortex and memory processed by the limbic systems (*16, 17*).

One of distinct targets of EN, the ventral CA1 (vCA1), is known to coordinate in a range of behaviors related to exploration and recognition memory, and these functions are thought to be regulated by its afferent inputs in a domain-specific manner (*18–26*). We, therefore, hypothesized that EN is a key node for establishment of recognition memory. To address this, we set out to characterize circuit and function of EN neurons defined by their projection to vCA1 (EN ^vCA1-proj.^ neurons) using genetic tools and mouse models of social and object recognition memory.

We found EN ^vCA1-proj.^ neurons innervated multiple components of the limbic system except amygdala and prefrontal cortex and produced potent feedforward inhibitory control over vCA1 pyramidal neurons. During recognition memory test, activity of EN ^vCA1-proj.^ neurons were condensed around conspecifics or objects where mice spent most time on. However, disruption of EN ^vCA1-proj.^ activity only impaired memory-guided exploration of novel stimuli without affecting innate exploration induced by novelty. These findings demonstrate that EN subserves some of the key functions required for recognition memory governed by specific limbic system.

## Results

### Endopiriform represents a major afferent of the ventral CA1

To assess the significance of EN as a vCA1 afferents, we injected a retrograde tracer (60 nl) into the vCA1 of mice and compared the number of the retrogradely labeled neurons in the EN and other brain regions (**Fig. 1A, B, S1A**, **Methods**). We found labeled neurons in multiple areas including the septum, entorhinal cortex, and basolateral amygdala (**Fig. 1C-F and Fig. S1B**) (*27, 28*). In addition to these well-known vCA1 afferents, prominent labeling was observed in the area defined as EN in Allen Brain Atlas (**Fig. 1G, Methods**). The number of presynaptic neurons in EN was ∼10 fold lower than entorhinal cortex (data not shown). However, the normalized count of presynaptic neurons in EN was significantly greater than basolateral amygdala, lateral septum, and medial septum (**Fig. 1H**).

**Figure 1.**
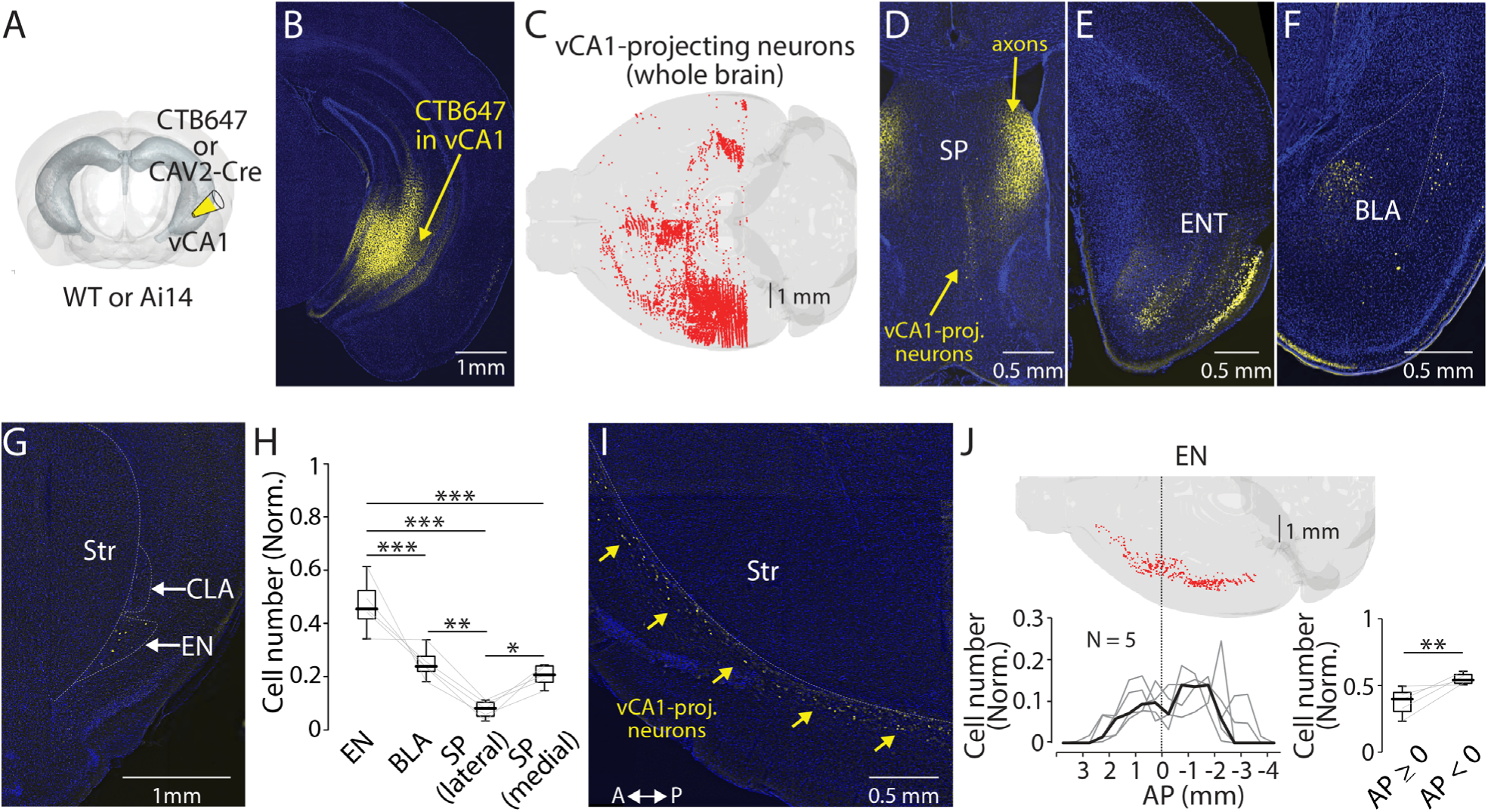
Endopiriform projects to the ventral CA1. (**A**) Schematic diagram of a retrograde tracer injection performed. CAV-2-Cre or CTB647 was used in combination with Ai14 or wild-type mice, respectively. (**B**) An example fluorescent image of injection site. (**C**) Distribution of retrogradely labeled vCA1-projecting neurons in entire brain as a result of injection in panel B. (**D-G**) Fluorescent images showing retrogradely labeled vCA1-projecitng neurons in septum (SP), entorhinal cortex (ENT), basolateral amygdala (BLA), and endopiriform (EN). Str: Striatum, CLA: Claustrum. (**H**) Box plots showing relative number of vCA1-projecting neurons in different brain areas. Cell number in each brain area were normalized by summed cell number of EN, BLA, SP (lateral), and SP (medial). Cell numbers in each brain areas of individual brains are presented in **Table 1**. Median value for EN, BLA, SP (lateral), and SP (medial) is 0.45, 0.24, 0.08, and 0.21, respectively. Statistics: EN vs BLA, SP (lateral), or SP (medial), p < 0.001 for all; BLA vs SP (lateral) or SP (medial): p = 0.002 and 0.694, respectively; SP (lateral) vs SP (medial) p = 0.023. One-way ANOVA, repeated comparison with Tukey-Kramer test. N = 5. (**I**) An example fluorescent image of horizontal section showing retrogradely labeled vCA1-projecting neurons in EN. A: anterior, P: posterior. (**J**) Top: Example map showing distribution of vCA1-projecting neurons in entire EN in one mouse. Bottom left: A plot showing a quantified distribution at bin size of 1 mm. Each bin was normalized to total number of labeled neurons in EN. Data from each mouse is shown in grey. Median is indicated in black. Bottom right: Box plots comparing a number of vCA1-projecting neurons in anterior and posterior EN. Data were normalized to a total number of labeled neurons in EN. Median value for anterior vs posterior was 0.39 vs 0.54. Statistics: p = 0.008, Rank-sum.

**Table 1.** Related to Fig. 1H.

| Brain areas | Brain #1 | Brain #2 | Brain #3 | Brain #4 | Brain #5 |
| --- | --- | --- | --- | --- | --- |
| EN | 288 | 201 | 310 | 210 | 250 |
| BLA | 85 | 107 | 175 | 208 | 118 |
| SP (lateral) | 27 | 36 | 69 | 69 | 17 |
| SP (medial) | 69 | 111 | 130 | 127 | 121 |
| **Sum** | **469** | **455** | **684** | **614** | **506** |

The vCA1-projecting neurons in the EN were spread along antero-posterior axis with two peak densities, one in anterior and another in posterior to the bregma (**Fig. 1I, J**). Number of labeled neurons was greater in posterior EN (**Fig. 1J**). In contrast, the same analysis along dorso-ventral axis showed a single dominant peak at the depth consistent with the location of EN (**Fig. S1C**). To further confirm the location of these vCA1-projecting neurons, we injected a retrograde tracer (60 nl for each) in one color into vCA1 and a retrograde tracer in another color into a cortical area known to receive projections from EN or “CLA” representing a dorsal part of CLA complex (**Fig. 2A**). We targeted the prefrontal cortex to label EN, and the anterior cingulate cortex, the motor cortex, and the dorsomedial entorhinal cortex to label CLA (**Fig. 2C-F**) (*29–32*).

**Figure 2.**
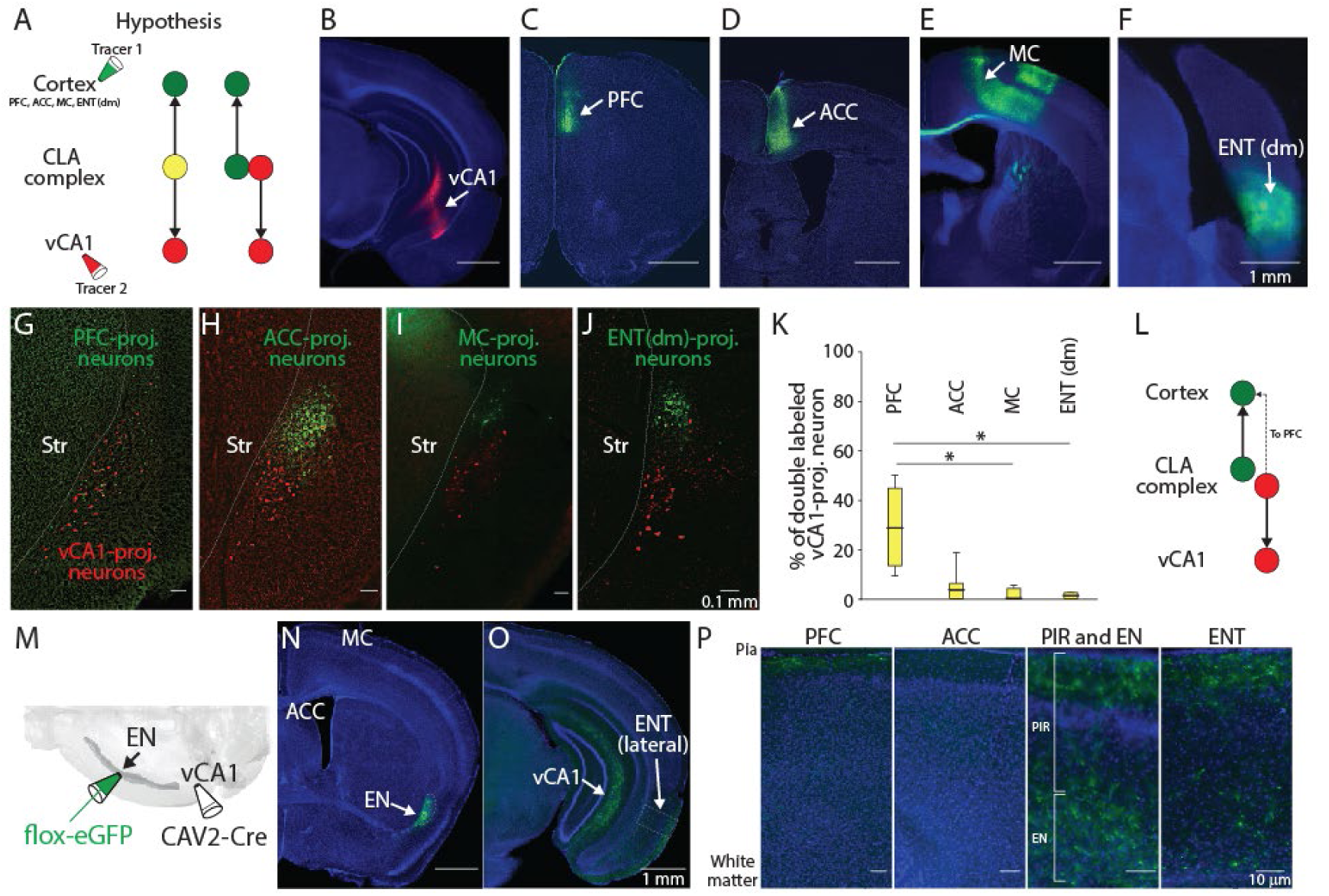
Axon branching of vCA1-projecting EN neurons. (**A**) Schematic diagram of injection performed and two hypothesized outcomes. Yellow color indicates double-labeled neurons. PFC: Prefrontal cortex, ACC: Anterior cingulate cortex, MC: Motor cortex, ENT(dm): Dorsomedial entorhinal cortex, CLA: claustrum (**B-F**) Example fluorescent images of injection site for each region indicated. (**G-J**) Example fluorescent images of retrogradely labeled neurons in CLA complex. vCA1-projecting neurons were colored in red and cortex-projecting neurons were colored in green. (**K**) Box plots showing percentage of double-labeled vCA1-projecting neurons for each cortical injection. Medial value for PFC, ACC, MC, and ENT(dm) was 28.6, 4.6, 0, and 1.2, respectively. Statistics: PFC vs ACC, MC or ENT(dm), p = 0.405, 0.015 and 0.047, respectively. ACC vs MC or ENT (dm), p = 0.422 and 0.64, respectively. MC vs ENT (dm), p = 0.993. One-way ANOVA with post-hoc Kruskal-Wallis test. Slice and animal number: 16 and 4 for PFC, 20 and 5 for ACC, 10 and 3 for MC, and 18 and 4 for ENT(dm). (**L**) Schematic diagram of projection pattern found in the experiment shown in A. (**M**) Schematic diagram of the intersectional approach performed to label vCA1-projecting EN neurons. (**N-O**) Example fluorescent images showing vCA1-projectig axons in different brain areas. All images were from the same mouse. (**P**) Magnified images showing vCA1-projecting axons in cortical layers. Magnified areas are indicated in **O** and fig. **S2A-C** as dashed squares. PIR: Piriform cortex

We found there is a major spatial overlap in labeling of vCA1-projecting neurons with prefrontal cortex-projecting neurons, but not with other cortical projection neurons (**Fig. 2G-J**). Moreover, around 30% of vCA1-projecting neurons were found to be double-labelled when a cortical injection was made into the prefrontal cortex, suggesting some vCA1-projecting neurons send axons to this cortical area (**Fig. 2K, L**).

To further determine the cortical and subcortical innervation pattern of vCA1-projecting EN neurons, we used an intersectional approach to label their somata and axons with eGFP. CAV2-Cre (60 nl) was injected into the vCA1 and flox-eGFP (100 nl) was injected into the EN (**Fig. 2M**). Apart from vCA1, axons from vCA1-projecting neurons were found in various cortical areas including prefrontal cortex, lateral entorhinal cortex, and piriform cortex (**Fig. 2N-P and Fig. S2A-C**). Projection to prefrontal cortex was much sparser relative to that of vCA1, as expected based on the retrograde labeling data (**Fig. 2K**). The connectivity of vCA1-projecting EN neurons to the amygdala, which represents the major component of limbic systems, was also sparse relative to vCA1 (**Fig. S2C**), indicating vCA1 is the main target of these EN neurons in this system. Within the hippocampus, very sparse EN axons were observed in vCA3 and dentate gyrus (**Fig. S2D**).

Taken together, these findings indicate EN represents a major afferent of vCA1. Moreover, EN neurons projecting their axons to vCA1 also send strong collaterals to lateral entorhinal cortex and piriform cortex, but relatively weak to the prefrontal cortex or other limbic structures. Since our subsequent study focused on EN neurons defined by their projection to vCA1, we refer to them as EN^vCA1-proj.^ neurons.

### Projection pattern and synaptic connectivity of EN axons in vCA1

We next investigated projection pattern of EN axons in hippocampus by injecting an anterograde tracer (AAV-GFP, 60 nl) into the EN (**Fig. 3A, B**). Analysis of different parts of hippocampal section indicated EN axons were mostly confined in SLM layer of distal part of vCA1 (**Fig. 3C, D**). Projection to intermediate or dorsal CA1 was limited or undetectable (**Fig. S3A-C**). Since SLM mainly consists of GABAergic neurons (*33*), we hypothesized that EN axons form synapses with this cell type. To test this, we first applied monosynaptic rabies tracing technique to Vgat-Cre mice (**Fig. 4A, Methods**). The distribution of starter cells (identified by GFP and mCherry co-labeling) was confirmed to be within the ventral-intermediate region of hippocampus using AMaSiNe (**Fig. 4B, S4A**) (*34*). In these mice, the presynaptic neurons were consistently observed in EN in addition to other expected areas (e.g., septum and entorhinal cortex) (**Fig. 4C** and **D, S4B-F**) (*35, 36*). The density of the presynaptic neurons in EN was greater at the posterior region to the bregma (**Fig. 4D**), consistent with our earlier observation (**Fig. 1J**).

**Figure 3.**
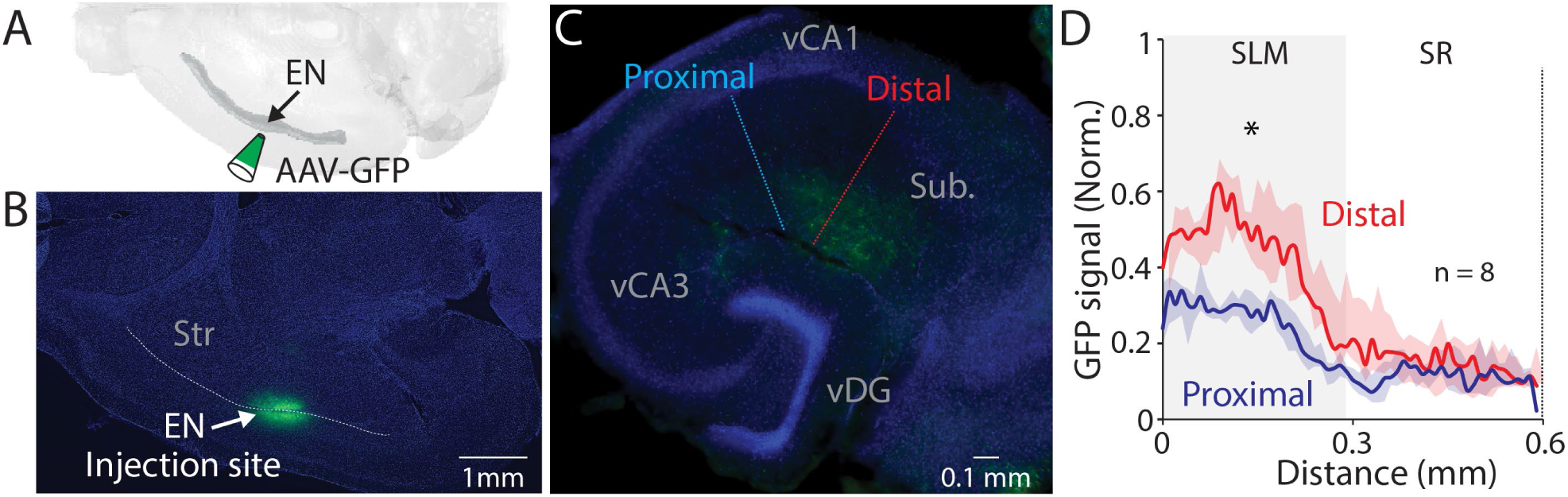
EN projection pattern in vCA1. (**A**) Schematic diagram of injection performed. (**B**) An example fluorescent image of horizontal brain section containing injection site in EN. (**C**) An example fluorescent image of ventral hippocampal slice with EN axons resulted from the injection in panel B. Dashed lines indicate the distal and proximal vCA1 regions where GFP signals were measured for the plot in panel D. (**D**) Normalized GFP signals across the layer. All signals were normalized by maximum GFP signal in the distal subregion. Median of area under the curve: distal; 11.12, proximal; 5.99. distal vs proximal, p = 0.016. Wilcoxon signed-rank test. n = 8 slices (N = 4 mice). SR: stratum radiatum, Py: pyramidal layer.

**Figure 4.**
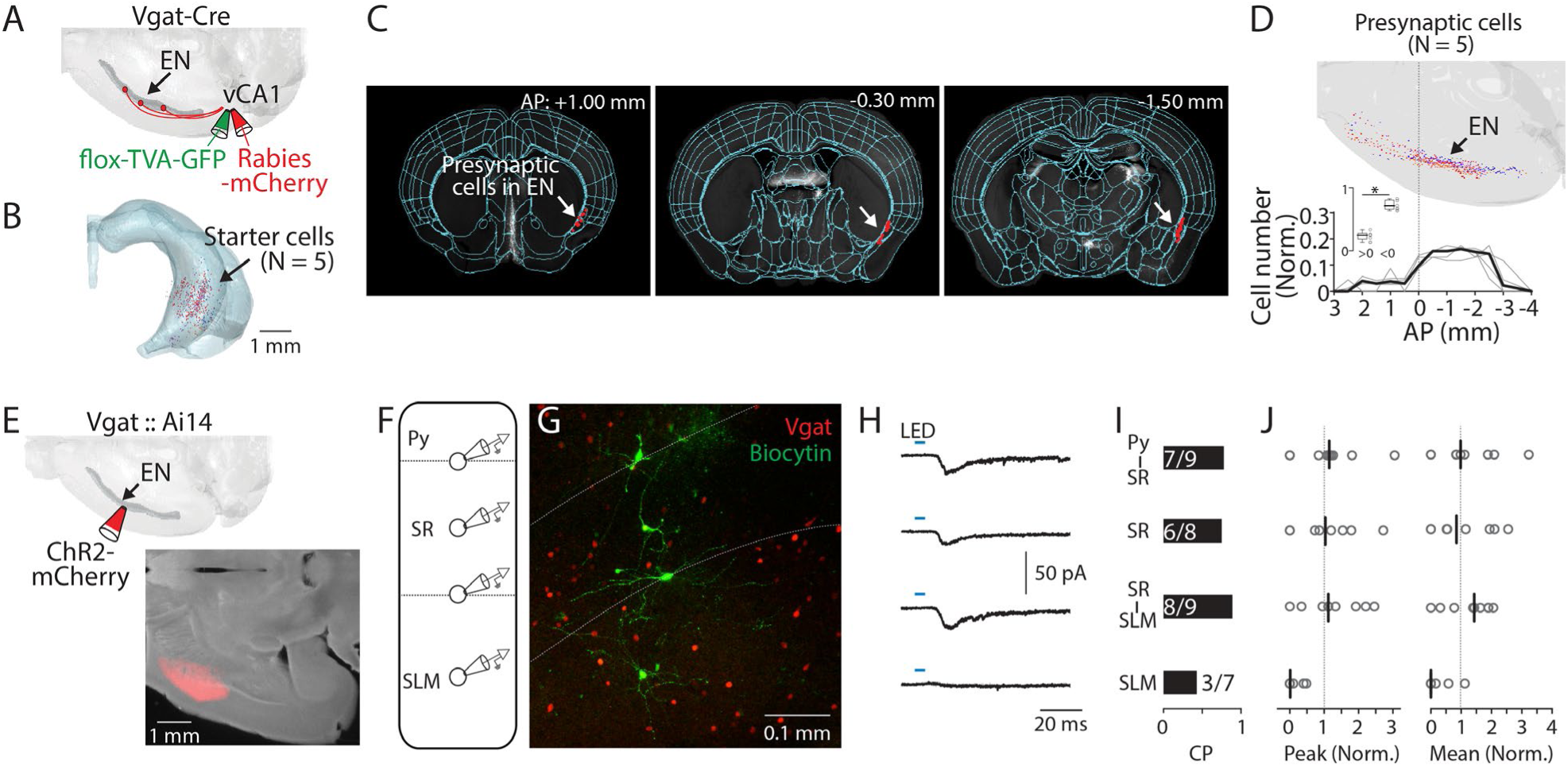
EN^vCA1-proj.^ neurons target vCA1 interneurons. (**A**) Schematic diagram of injection performed for monosynaptic rabies tracing. Flox-TVA-GFP (60 nl) was injected into the vCA1. 4 weeks later, Rabies-mCherry (60 nl) was injected. (**B**) Location of starter cells in the hippocampus. Data from 5 brains (each brain represented by different color) were overlaid. (**C**) Example images showing presynaptic cells in EN (white arrow). (**D**) Top: Location of presynaptic EN neurons. Data from 5 brains were overlaid (different brains were coded by different colors). Bottom: Plots indicate distribution of presynaptic cells in EN along AP axis. Cell counts in a given bin (1 mm) were normalized by total cell number in EN. Thin gray lines indicate individual data. A thick black line indicates median. Inset showing plot comparing cell number in anterior and posterior location as in Fig. 1J. Median: anterior vs posterior, 0.25 vs 0.72, p = 0.031; Wilcoxon signed-rank test. (**E**) Top: Schematic diagram of injection performed. Bottom: An example fluorescent image of injection site in EN. (**F**) Schematic diagram of recordings performed. Recordings were performed from identified GABAergic neurons sequentially in a random order. (**G**) An example confocal image of an acute vCA1 slice used for recordings. Recorded GABAergic neurons were filled with biocytin and post-labeled with Alexa488. White dashed lines indicate border of Py-SR and SR-SLM. (**H**) Median traces of photo-evoked EPSCs recorded from neurons at different laminar position. (**I**) Connection probability (CP) of EN axons and recorded neurons at different laminar positions. Numbers in bars indicate number of response positive neurons /total recorded neurons. (**J**) Left: Peak amplitudes of evoked EPSCs (normalized to total input from recorded slice). Right: the same as left but for mean amplitude. Gray circles indicate each neuron. Black lines indicate median. Median (peak amp): Py-SR vs SR, SR-SLM or SLM: 1.1 vs 1.0 vs 1.1 vs 0.0. [Py-SR vs SR, SR/SLM vs SLM], p = 0.999, 0.988, 0.076; [SR vs SR-SLM or SLM], p = 0.972, 0.111; [SR-SLM vs SLM], p = 0.039; one-way ANOVA, repeated comparison with Tukey-Kramer test. Median (mean amp): Py-SR vs SR, SR-SLM, or SLM: 1.0 vs 0.8 vs 1.4 vs 0.0. [Py-SR vs SR, SR-SLM or SLM], p = 0.991, 0.999, and 0.141; [SR vs SR-SLM or SLM], p = 0.978, and 0.251; [SR-SLM vs SLM], p = 0.114; one-way ANOVA, repeated comparison with Tukey-Kramer test. N = 5 mice.

To determine how EN inputs are integrated into the local circuit in vCA1, we systematically assessed the connections between EN axons and GABAergic neurons in different layers of distal vCA1. We expressed channelrhodopsin-2 (ChR2) into the EN axons by injecting AAV-ChR2-mCherry (100 nl) into EN and performed whole-cell recording from GABAergic neurons in acute vCA1 slices in the presence of TTX (1 µM) and 4-AP (100 µM) to isolate monosynaptic connection (*37*). To ensure the recording from GABAergic neurons, we used Vgat::Ai14 mice in which all GABAergic neurons are labeled with tdTomato (**Fig. 4E-G**). We recorded from 3 to 4 GABAergic neurons located in different laminar position in distal vCA1 (Py-SR border, SR, SR-SLM border, or SLM) in the same slice to construct a “laminar profile” of EN→vCA1 inputs (**Fig. 4F and G**). Whole-field LED stimulation evoked excitatory postsynaptic current (EPSC) in most GABAergic neurons recorded. However, contrary to the expectation, connection probability and strength were lowest in neurons in SLM compared to neurons in other layers (**Fig. 4H-J**). Taken together, these results indicate EN axons preferentially innervate the GABAergic neurons spanning across the Py-SR border to SR-SLM border in distal vCA1.

### EN axons produce feedforward inhibition onto vCA1 pyramidal neurons

SLM also contains the apical tuft dendrite of pyramidal neurons, which could be a target of EN axons in addition to GABAergic neurons. To test this, we expressed ChR2 in EN axons but made whole-cell recording from pyramidal neurons in distal vCA1 in the presence of TTX and 4-AP. Since CA1 pyramidal neurons can be differentiated into superficial and deep layer neurons with distinct anatomy and function (*38*), we recorded from neurons in each layer as well as nearby GABAergic neurons at the Py-SR and SR-SLM borders for the comparison (**Fig. 5A**). We found monosynaptic input to superficial and deep layer pyramidal neurons was similar, and monosynaptic input to pyramidal neurons (superficial and deep pooled) and GABAergic neurons was also comparable (**Fig. S5**).

**Figure 5.**
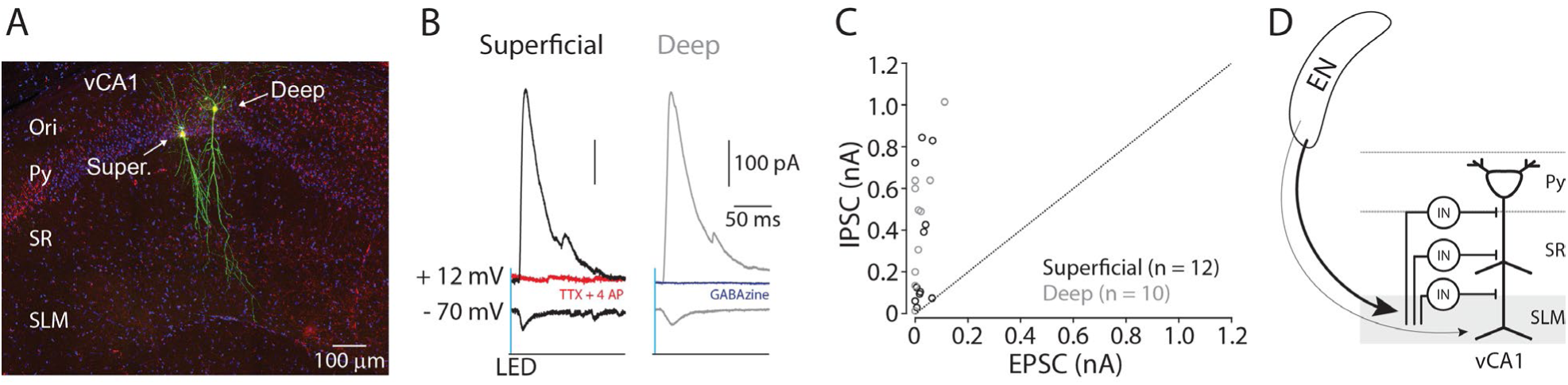
EN disynaptically inhibits vCA1 pyramidal neurons. (**A**) A confocal image of recorded pyramidal neurons in vCA1. Recorded pyramidal neurons were filled with biocytin and post-labelled with Alexa488. Ori.: oriens, Super.: superficial layer pyramidal neuron. Deep: deep layer pyramidal neuron. (**B**) Example traces of photo-evoked EPSCs and IPSCs in superficial (black trace) and deep (gray trace) pyramidal neurons. Red trace: after TTX and 4AP. Blue trace: after gabazine. (**C**) Scatter plots of peak amplitudes of EPSCs and IPSCs for each neuron. Dashed line indicates a unitary line. Superficial neurons: empty Black circles. Deep neurons: Gray circles. N = 4 mice. (**D**) A summary diagram of EN→vCA1 circuit. In vCA1, EN axons innervate GABAergic neurons (IN), which in turn inhibits pyramidal neurons (both superficial and deep).

Since EN axons innervated multiple GABAergic neurons across different layers, we hypothesized they may exert stronger inhibition onto pyramidal neurons through feed-forward inhibition. To test this, we recorded EPSC and inhibitory postsynaptic current (IPSC) from superficial and deep pyramidal neurons using cesium-based internal solution in the brain slices expressing ChR2 in the EN axons (**Methods**). At the command potential of −70 mV, photo-stimulation of EN axons evoked EPSCs in majority of pyramidal neurons in both layers (7 out of 10 neurons in superficial and 5 out of 10 neurons in deep) **(Fig. 5B**). At the command potential of +12 mV, the same stimulation evoked outward postsynaptic current in the same neurons, including those without EPSCs (10 out of 10 for superficial and deep) (**Fig. 5B**). This outward current was abolished by application of TTX and 4-AP or gabazine (10 µM), indicating it is disynaptically driven and mediated by inhibitory GABA_A_ receptor (**Fig. 5B**). When relative strength of EPSCs and IPSCs were compared for each neuron, the strength of IPSCs overwhelmed the EPSCs in all cases (**Fig. 5C**). This indicates EN axons disynaptically inhibit pyramidal neurons in vCA1 (**Fig. 5D**).

### EN^vCA1-proj.^ neurons receive inputs from the piriform cortex

We next examined afferents that may drive the EN→vCA1 circuit. To address this, retrograde CAV2-Cre was injected into vCA1 to express Cre into EN^vCA1-proj.^ neurons, then applied monosynaptic rabies tracing technique to specifically label its presynaptic neurons in wild type mice (**Fig. 6A**). Starter cells and presynaptic neurons were spatially mapped and quantified using AMaSiNe (**Fig. 6B and C, S6**). Since presynaptic neurons were sparse in contralateral hemisphere, data from ipsilateral hemisphere was analyzed. Moreover, we applied minimum labeling threshold to determine the brain region for further analysis (**Methods**). We found the piriform cortex contained highest number of presynaptic neurons followed by EN (**Fig. 6D-F**). Within piriform cortex, layer 2 neurons were the major source of afferent (**Fig. 6G**). These data indicate EN^vCA1-proj.^ neurons receive major input from piriform cortex and form a recurrent connection with neurons in EN (**Fig. 6H**).

**Figure 6.**
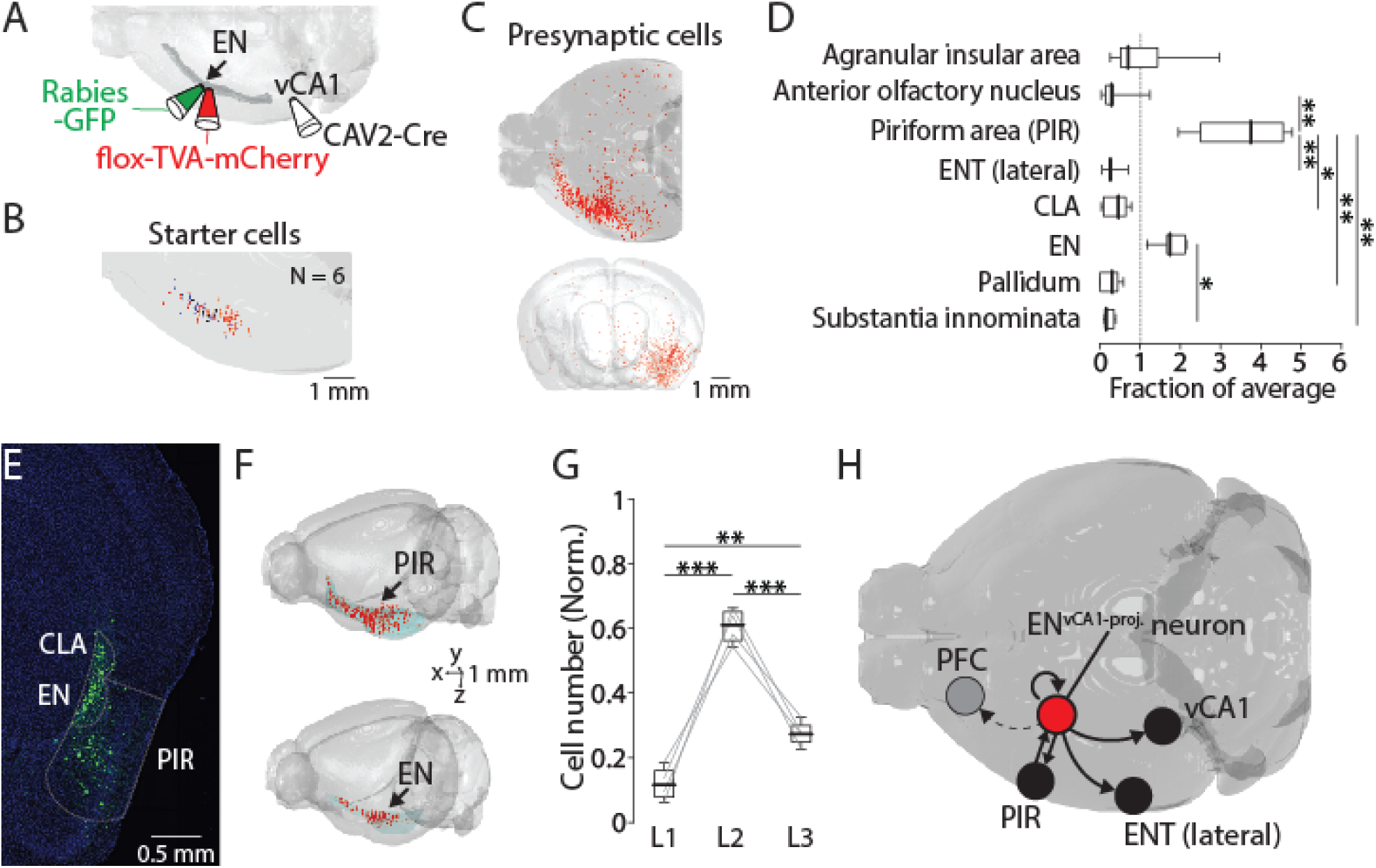
EN^vCA1-proj.^ neurons receive inputs from the piriform cortex and within EN. (**A**) Schematic diagram of performed. After CAV2-Cre (60 nl) injection into the vCA1, flox-TVA-mCherry (100 nl) was injected into the EN. 4 weeks later, Rabies-GFP (100 nl) was injected into the EN. (**B**) Location of starter cells in EN. Data from 6 brains were overlaid. (**C**) An example map of presynaptic cells in one brain. Top: Horizontal view. Injections were made in left hemisphere. Bottom: Front view. (**D**) Number of presynaptic cells in different brain areas (normalized to total presynaptic cell number). Median for AIA, AON, PIR, ENT(lateral), CLA, EN, Pallidum, and SI (in fraction): 0.7, 0.3, 3.8, 0.3, 0.5, 1.7, 0.3, and 0.2. [AON, ENT(lateral), CLA, Pallidum, SI] vs PIR: p = 0.016, 0.008, 0.048, 0.006, and 0.003. EN vs SI, p = 0.043. one-way ANOVA with post-hoc Kruskal-Wallis test. Full list of statistical tests and p values are in **Table 2**. (**E**) An example fluorescent image of coronal slice containing EN and PIR and labeled presynaptic cells. (**F**) Example maps of presynaptic cells in PIR (top) and EN (bottom). (**G**) Number of presynaptic cells in PIR in different layers (normalized to the total number of labeled neurons in PIR). Median: L1 vs L2 vs L3, 0.11 vs 0.61 vs 0.27. L1 vs L2, p < 0.001; L1 vs L3, p = 0.005; L2 vs L3, p < 0.001; one-way ANOVA, repeated comparison with Tukey-Kramer test. (**H**) Summary diagram showing input-output circuit of EN^vCA1-proj.^ neurons (red circle).

**Table 2.** Related to Fig.6D.

| Compared brain areas |  | p value |
| --- | --- | --- |
| Agranular insular | AOB | 0.764 |
| Agranular insular | PIR | 0.617 |
| Agranular insular | ENT | 0.624 |
| Agranular insular | CLA | 0.921 |
| Agranular insular | EN | 0.971 |
| Agranular insular | Palidum | 0.588 |
| Agranular insular | Substantia innominata | 0.44 |
| AOB | PIR | 0.016 |
| AOB | ENT | 1 |
| AOB | CLA | 1 |
| AOB | EN | 0.156 |
| AOB | Palidum | 1 |
| AOB | Substantia innominata | 1 |
| PIR | ENT | 0.008 |
| PIR | CLA | 0.048 |
| PIR | EN | 0.994 |
| PIR | Palidum | 0.006 |
| PIR | Substantia innominata | 0.003 |
| ENT | CLA | 1 |
| ENT | EN | 0.091 |
| ENT | Palidum | 1 |
| ENT | Substantia innominata | 1 |
| CLA | EN | 0.317 |
| CLA | Palidum | 0.999 |
| CLA | Substantia innominata | 0.992 |
| EN | Palidum | 0.079 |
| EN | Substantia innominata | 0.043 |
| Palidum | Substantia innominata | 1 |

### The activity of EN^vCA1-proj.^ neurons is correlated with the time mice spend in space

The downstream target of EN, vCA1, is implicated in social, odor and object recognition memory (*20–22*). To determine whether and how the activity of EN neurons is related to this function, we expressed GCaMP8s in EN^vCA1-proj.^ neurons and monitored their activity using fiber photometry while video recording mice performing social recognition memory test consisting of pretest, sociability test, and discrimination test (**Fig. 7A, B, Methods**). The nose position of the subject mouse was tracked during tests using DeepLabCut (**Fig. 7C**) (*39, 40*), and cumulative time and calcium event were mapped onto the arena space (**Fig 7C, S7**). We found these values were correlated in space across all sessions (i.e., the more time the mice spent in a given space, the more the EN^vCA1-proj.^ activity), including in the open field with no object or in the test arena with conspecifics or objects (**Fig. S7, S8A-B**).

**Figure 7.**
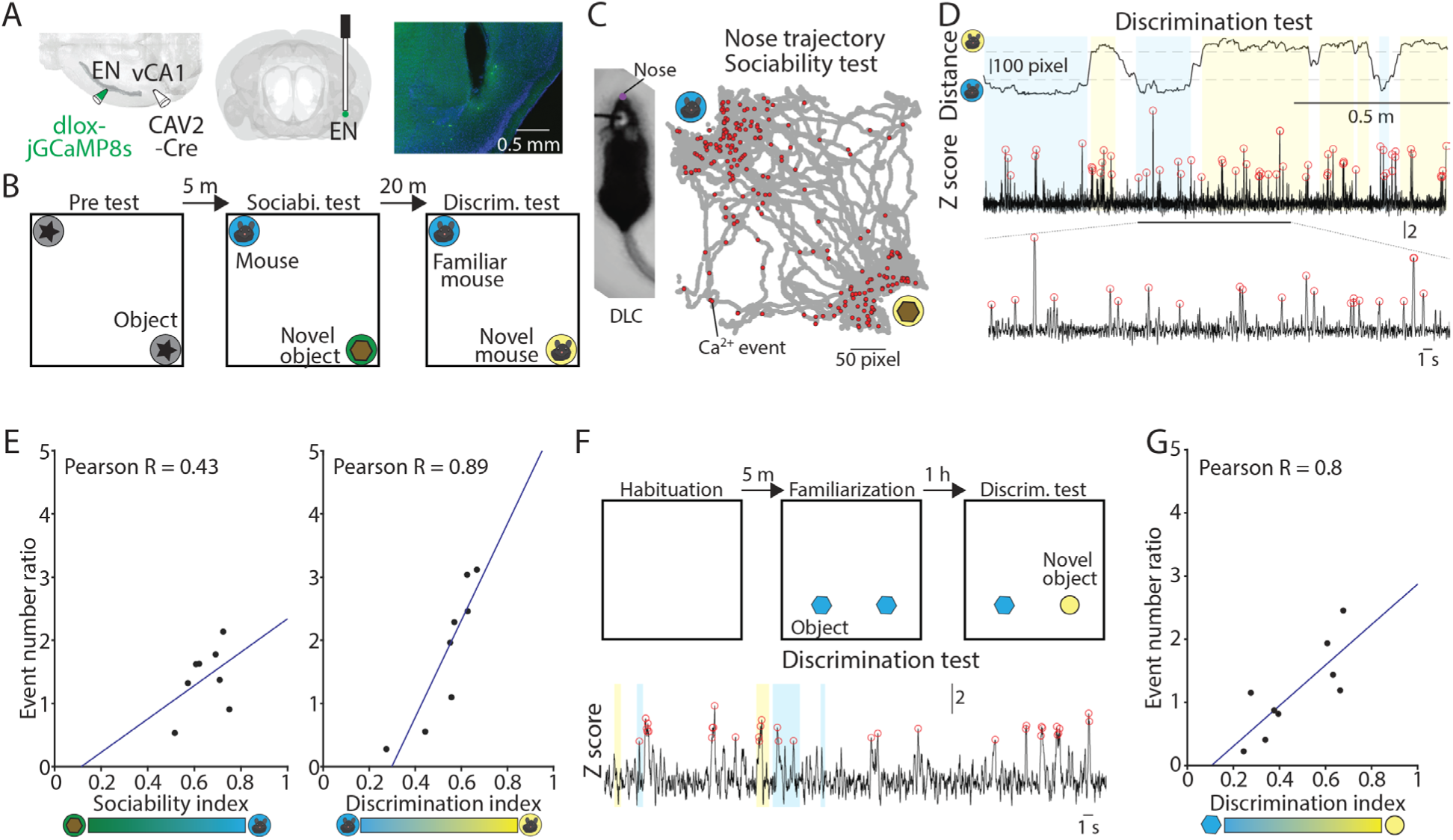
Activity of EN^vCA1-proj.^ neurons correlates with social/object discrimination performance. (**A**) Right and Middle: Schematic diagrams of injection and optic cannula implantation performed. CAV2-Cre (60 nl) was injected into the vCA1, and dlox-jGCaMP8s (100 nl) was injected into the EN. Left: An example image showing jGCaMP8s expressing EN^vCA1-proj.^ neurons and a tract of optic cannula. (**B**) The diagram of social recognition memory test. A test consisted of pretest (5 min), sociability test (10 min), and discrimination test (5 min). (**C**) Left: An example image of a mouse with nose point marker tracked by DLC. Right: An example map of nose points and calcium events in the arena during sociability test. (**D**) Nose distance vector (distance between subject’s nose point and a center point of novel/familiar mouse chambers) and simultaneously acquired calcium signals during discrimination test. Gray dashed lines indicate the border of interaction zones. Open red circles indicate the calcium event detected (threshold: Z > 2.58). A part of calcium signal trace was expanded at the bottom for clarity. (**E**) Left: Correlation between calcium event ratios (calcium event number in the mouse interaction zone/ calcium event number in the object interaction zone) and sociability index. p = 0.29, N = 8 mice. Right: same as in left panel, but for calcium event ratios (calcium event number in the unfamiliar mouse interaction zone/ calcium event number in the familiar mouse interaction zone) and discrimination index. p = 0.003, N = 8. (**F**) The diagram of novel object recognition memory test. A test consisted of habituation (5 min), familiarization (10 min), and discrimination test (5 min). (**G**) Same as in (**E**), but for calcium event number ratios (calcium event number in the unfamiliar object interaction zone/calcium event number in the familiar object interaction zone and discrimination index. p = 0.01. N = 9 mice.

Cumulative calcium events of EN^vCA1-proj.^ neurons became high towards the space around objects or conspecifics when they are present and was correlated with interaction times except for sociability session (**Fig. S8C-D**). In the sociability session, the cumulative calcium events around a novel conspecific and object were similar despite mice spending more time on conspecifics (**Fig. 7E left, S8C**). In contrast, in the social discrimination test, both the cumulative time and cumulative EN^vCA1-proj.^ activity were higher for an unfamiliar than familiar conspecific (**Fig. 7D-E, S8C**). As a result, EN^vCA1-proj.^ activity and behavior had a greater degree of correlation in social discrimination than sociability.

We also tested if a similar correlation occurs in a non-social context by recording EN^vCA1-proj.^ activity and behavior in the object discrimination test (**Fig. 7F**). The cumulative time and calcium events were significantly correlated across different sessions i.e., familiarization, and object discrimination (**Fig. 7G, S8E**), largely consistent with the findings in the social discrimination test.

These data indicate that the activity of EN^vCA1-proj.^ neurons is generally correlated with the time the mouse spends in a particular location at the basal level (i.e. open field) or task conditions. However, the degree of correlation appears to vary depending on the session, such that it was highest in the pretest/familiarization and social/object discrimination test, but lowest in the sociability test.

### Inhibition of EN^vCA1-proj.^ neurons impairs social/object recognition memory

Although EN^vCA1-proj.^ activity was correlated with the behavior in pretest/familiarization and discrimination test better than sociability test, their causality is unclear. To address this, we inhibited the activity of EN^vCA1-proj.^ neurons using inhibitory DREADD (hM4Di) during social and object discrimination tests (**Fig. 8A-C and h, S9A**) (*41*). To specifically express hM4Di in EN^vCA1-proj.^ neurons, CAV-2-Cre (60 nl) and AAV-flox-hM4Di (100 nl for each coordinate) were bilaterally injected into vCA1 and EN, respectively (**Fig. 8A-B**). hM4Di was replaced with tdTomato for control group. We made a protocol that allows within-subject comparison, such that a mouse goes through a test with chlozapine-N-oxide (CNO, 1 mg/ kg) treatment then the same test with saline treatment the next day, or vice versa (**Fig. 8C and H**). In social discrimination test, CNO treatment did not affect the pretest or sociability in hM4Di or control group, but specifically impaired social discrimination in hM4Di group (**Fig. 8D-G, S9B-G**). A similar effect was also observed in novel object discrimination (**Fig. 8I, S9D**).

**Figure 8.**
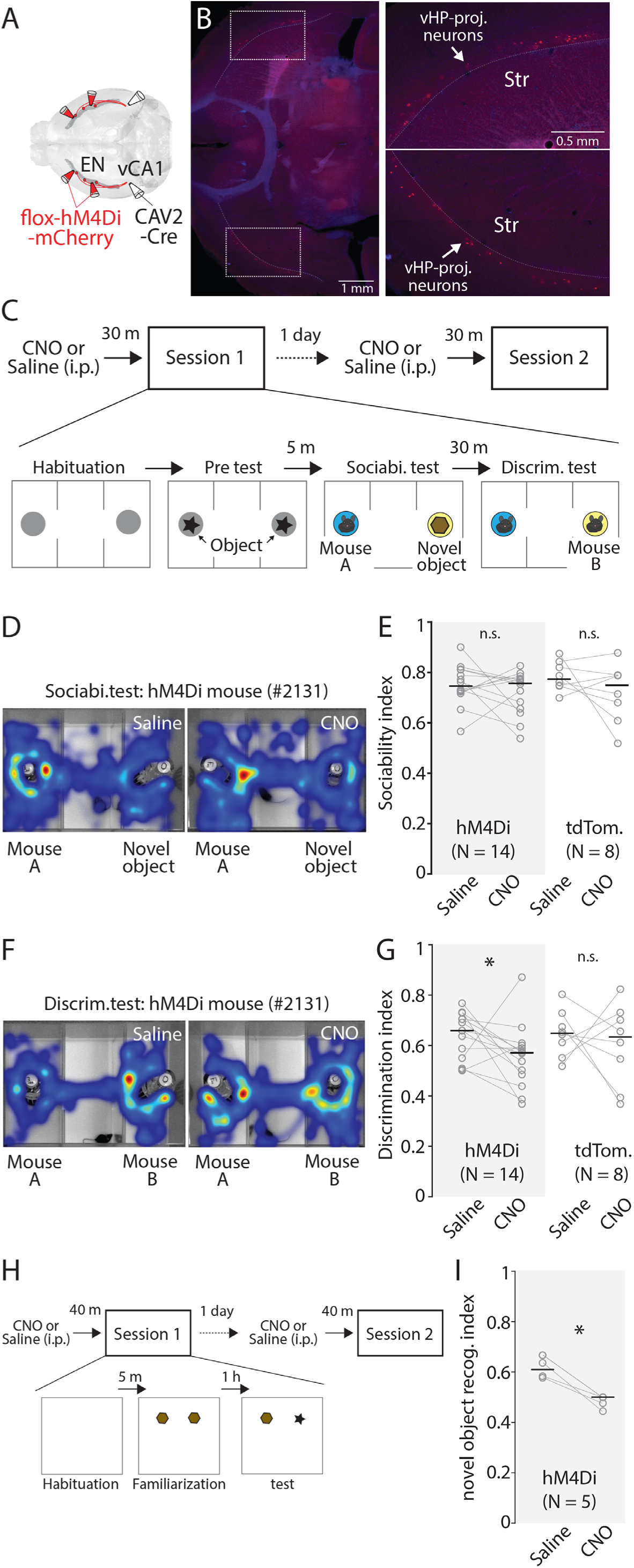
Chemogenetic inhibition of EN^vCA1-proj.^ neurons specifically impairs recognition memory. (**A**) Schematic diagram of injection performed. (**B**) Left: An example fluorescent image showing expression of hM4Di in EN^CA1-proj.^ neurons. Right: Expansion of areas marked by white rectangle in image on the left. (**C**) The diagram of social recognition memory test. A session consisted of habituation (5 min), Pretest (5 min), Sociability test (10 min), and discrimination test (5 min). The group treated with a saline in session 1 was treated with CNO (1mg/ kg) in session 2 the following day. (**D**) Example cumulative times heat map in arena during sociability tests for the same mouse under two different treatments. **(E)** Plots comparing sociability index for hM4Di and tdTomato mice under two different treatments. Gray empty circles indicate each mouse. Gray thin lines show paired data from one mouse. Thick black bars indicate median. Median sociability index: hM4Di mice (Saline vs CNO), 0.75 vs 0.76, p = 0.391; signed-rank test; tdTomato mice (Saline vs CNO), 0.77 vs 0.75, p = 0.383; signed-rank test. (**F**) Example cumulative time heat maps in arena during discrimination tests for the same mouse under two different treatments. (**G**) Comparison of discrimination index for hM4Di and tdTomato mice under two different treatments. Gray empty circles indicate each mouse. Gray thin lines show paired data from one mouse. Thick black bars indicate median. Median discrimination index: hM4Di mice (Saline vs CNO), 0.66 vs 0.57, p = 0.042; signed-rank test; tdTomato mice (Saline vs CNO), 0.65 vs 0.63, p = 0.742; signed-rank test. (**H**) The diagram of novel object recognition memory test. A session consisted of habituation (5 min), familiarization (10 min), and discrimination test (5 min). CNO treatment was the same as in **C**. (**I**) Same as in **G**, but for novel object discrimination index. Median discrimination index: hM4Di mice (Saline vs CNO), 0.64 vs 0.5, p = 0.003; signed-rank test.

Apart from recognition memory, vCA1 is implicated in anxiety and associative fear memory (*42, 43*), thus, we tested contribution of EN^vCA1-proj.^ neurons to these functions. The anxiety level as determined by locomotor activity (*44*) and sociability was not affected by CNO treatment in hM4Di group (**Fig. 8E, S9E**). Similarly, fear memory as assessed by the freezing response during training (with trace fear conditioning) or recall to context or tone was not affected by CNO treatment in hM4Di group (**Fig. S10**).

Taken together, these data indicate that EN^vCA1-proj.^ activity is not causally related to innate exploration behavior induced by novelty, or anxiety or fear memory. However, their activity is necessary for mice to discriminate between familiar and unfamiliar conspecific or object, suggesting their major role in general recognition memory.

## Discussion

We characterized the circuit and function of EN neurons targeting specific subregions and layers of vCA1. In contrast to other parts of CLA complex, EN^vCA1-proj.^ neurons were reciprocally connected with the piriform cortex representing a major downstream target of the olfactory bulb (*45*). Since social odor information is crucial for discriminating conspecifics in rodents, this circuit motif may predict the predominant role of EN^vCA1-proj.^ neurons in social recognition memory, given that social odor can engage multiple olfactory pathways innervating the piriform cortex (*46*). However, we found that these neurons contributed to broader domains of recognition memory. This is supported by the observations that the inhibition of EN^vCA1-proj.^ activity impaired both social and non-social discrimination.

In all phases of recognition memory test, EN ^vCA1-proj.^ activity was highly correlated with the time mice spent in given locations and this correlation was biased towards novel stimuli when mice were engaged in their exploration irrespective of the stimuli being social or non-social (**Fig. 9A**). Taken together, these data suggest that the function of EN^vCA1-proj.^ neurons was primarily related to the detection of and attention to novel stimuli. The recurrent connections within EN, which is another prominent feature of EN^vCA1-proj.^ neurons’ network, potentially support these processes to help establish the recognition memory (*47*).

**Figure 9.**
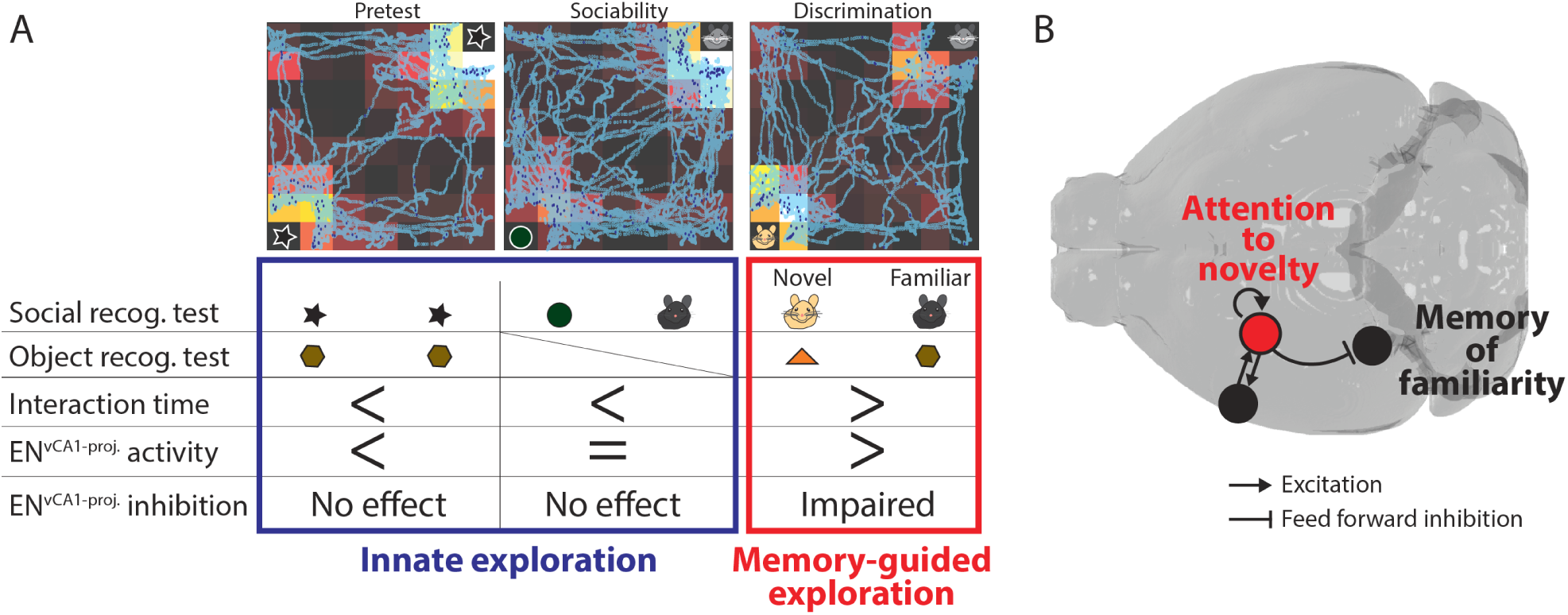
The role of EN^vCA1-proj.^ neurons in recognition memory. **(A)** Summary of findings from *in vivo* experiments. **(B)** The model of EN^vCA1-proj.^ circuit function in recognition memory. During memory-guided exploration, recurrent circuits in EN^vCA1-proj.^ neurons maintain the attentional response to novel stimuli while regulating the degree of response to familiar stimuli in vCA1 via feedforward inhibition.

Attentional processes mediated by EN^vCA1-proj.^ neurons can be considered of particular relevance for memory-guided behavior, given that inhibition of EN^vCA1-proj.^ activity selectively impaired conspecific/object discrimination but not innate exploratory behavior provoked by novelty as seen in pretest, sociability, and familiarization (**Fig. 9A**). The correlation of EN^vCA1-proj.^ activity with novel object preference in the pretest nevertheless suggests that these neurons ‘represent’ the innate preference without driving it. This functional specialization, likely associated with its unique circuit connectivity to the limbic system, could differentiate EN from CLA.

Studies on EN are generally scarce, and the available evidence of EN projections to vCA1 suggests that these projections are sparse (*14*). Potential reasons include the difficulty in clearly delineating the EN from CLA complex and non-uniform distribution of EN^vCA1-proj.^ neurons along antero-posterior axis. By leveraging contemporary circuit analysis tools, we demonstrated that the EN→vCA1 circuit is non-trivial; EN^vCA1-proj.^ neurons were more numerous than commonly studied vCA1 afferents, including those from the septum and amygdala. Moreover, these axons exhibited a potent disynaptic inhibition of distal vCA1 pyramidal neurons, indicating their major functional implications for vCA1 activity.

Our data indicated that although EN axons terminated at the SLM of distal vCA1, they exhibited stronger connections to GABAergic neurons spanning from the Py-SR border to the SR-SLM border than to SLM neurons. The detection of EN inputs from GABAergic neurons at the Py-SR border was surprising, considering the distance between soma position and the EN axons. However, there are several GABAergic cell types below SLM that extend their primary dendrites into SLM (*48, 49*), likely explaining the observed connectivity pattern. Among these are chandelier cells found at the Py-SR border, which preferentially inhibits the axon initial segments of pyramidal neurons, and are thus well suited to exert the powerful feedforward inhibition of vCA1 pyramidal neurons observed in our recordings (*50*).

There are several potential circuit effects by which EN inputs could affect vCA1 function. One is the induction of non-linear events in vCA1 pyramidal neurons by acting in concert with other afferents (e.g., lateral entorhinal cortex targeting SLM) (*36*) to generate a specific output pattern in pyramidal assemblies representing task-relevant stimuli. Alternatively, EN inputs may improve signal to noise ratio of the information conveyed through the lateral entorhinal cortex, CA2 (in the case of social context), and possibly other afferents (e.g., CA3) (*51*). Another scenario is tuning of vCA1 pyramidal neurons’ response to familiar stimuli through dynamic combination of a weak monosynaptic excitatory input and strong disynaptic inhibitory input. Such mechanism suggests that at a milliseconds scale novelty (EN) and familiarity (vCA1) recognition might occur though alternating rather than simultaneous activity of brain microcircuits. Taken together, we propose a model for the role of EN^vCA1-proj.^ neurons in recognition memory by balancing memory-guided attentional responses to familiarity and novelty through combination of feedforward inhibition of vCA1 (a node for recognition of familiarity) and the recurrent circuits that contribute to sustain attention to novelty (**Fig. 9B**).

The model of promoting novelty detection by suppressing familiarity responses is consistent with previous observations showing that vCA1 pyramidal neurons predominantly respond to familiar stimuli whereas novelty response can only occur through activation of vCA1 interneurons (*52, 53*). Nevertheless, alternating activation of EN and vCA1 could prove essential for the behavioral relevance of EN activity associated with recognition memory, without affecting innate exploratory behavior to novelty. Accordingly, the latter behavior is primarily associated with the prefrontal cortex and amygdala, areas that are only weakly targeted by EN^vCA1-proj.^ neurons (**Fig. 2P, S2C**) (*54–56*).

In addition to advancing our understanding of the basic organization and function of brain circuits underlying higher cognitive processes, our findings suggest dysfunction of EN^vCA1-proj.^ neurons could be a key contributing factor to the deficits in CLA complex function and recognition memory found in neuropsychiatric disorders (*9, 10*). Discrete EN populations can thus emerge as pathophysiological substrate, but also as important treatment targets of key symptoms of these disorders.

## Materials and Methods

### Mice

All experiments were performed in accordance with standard ethical guidelines and were approved by the Danish national animal experiment committee (License number: 2021-15-0201-00801). Unless otherwise noted, all mice used were C57BL6/J mice or transgenic mice with C57BL6/J backgrounds. They were 2-4 months old at the time of the experiment. Transgenic mice used were B6.Cg-Gt (ROSA)26Sor^tm14(CAG-tdTomato)Hze^/J mice (Ai14, JAX007914) and B6J.129S6(FVB)-Slc32a1^tm2(cre)Lowl^/MwarJ mice (Vgat-Cre, JAX028862). Similar number of male and female mice were used for all experiments except for behavior tests. For behavior tests, male mice were used.

### Viruses and retrograde tracers

The information on viruses used in our experiments was the following. CAV-2-Cre (PVM); AAV1-mDlx-HBB-chl-dlox-TVA_2A_oG(rev)-dlox (v271-1, VVF); AAV1-hSyn1-dlox-TVA_2A_mCherry_2A_oG(rev)_dlox (v306-1, VVF); Rabies-GFP (NTNU viral core facility); Rabies-mCherry (NTNU viral core facility); AAV5-CAG-GFP (37525-AAV5, Addgene); AAV5-CAG.hChR2(H134R)-mCherry (10054-AAV5, Addgene); AAV8-hSyn-DIO-hM4Di (Gi) – mCherry (44362-AAV8, Addgene); AAV5-hEF1a-dlox-EGFP(rev)-dlox (v217-5, VVF); and retrograde AAVrg-CAG-GFP (37825-AAVrg, Addgene), AAV1-hSyn1-dlox-jGCaMP8s(rev) (v627-1, VVF). We also used retrograde traces of red Retrobeads (Lumafluor) and Cholera toxin subunit B conjugated with Alexa 647 (CTB647, Thermo Fisher).

### Stereotaxic injections

Stereotaxic injections were performed using a stereotaxic frame (Model 940, Kopf). Mice were anesthetized with isoflurane and were subcutaneously injected with buprenorphine (0.1 mg/kg) and metacam (1.5 mg/Kg) for post-operative pain relief. After incising the scalp over the cranium, a small hole was bored with a microdrill bit, and a beveled glass pipette (Wiretrol II, 5-000-2010, Drummond Scientific Company) back-filled with mineral oil and front-filled with the material to be injected was slowly inserted into a target coordinate. After injecting a small volume (50 - 100 nL) with a custom-made displacement injector (based on MO-10, Narishige), the pipette was left in place for 3 to 5 mins before slow retraction. The incision was closed with a nylon suture. The stereotaxic coordinates used were (relative to bregma, in mm): anteroposterior (AP) −3.0; mediolateral (ML) 3.12; dorsoventral (DV) 3.7 for ventral CA1; AP +2.0; ML 0.2; DV 1.5 and 2.5 for prefrontal cortex; AP 0.0; ML 0.2; DV 1.0 and 1.5 for anterior cingulate cortex; AP 0.0; ML 1.5; DV 0.3 and 0.7 for motor cortex; and AP −5.0; ML 3.0; DV 1.5 for dorsomedial entorhinal cortex. For endopiriform injection, we injected two sites from the following coordinates: AP −0.27, ML 3.2, DV 4.2; AP 0.0, ML 3.0, DV 4.2; or AP +1.0, ML 2.7, DV 4.0. Mice were thermally supported with a feedback-controlled heating pad maintained at ∼37 °C (ThermoStar Homeothermic system, RWD). Mice were used for experiments 3-5 weeks post injection.

### Implantation of the optic probe

After stereotaxic injection of CAV-2-Cre into the vCA1 and AAV1-hSyn1-dlox-jGCaMP8s into the EN (AP −0.27, ML 3.2, DV 4.2), a fiber optic cannula (400 μm core diameter, 4.5 mm length, NA 0.39, R-FOC-BF-400C-39NA, RWD) was inserted towards EN. Once in the correct depth, exposed brain surface around the implant was covered by Kwik-Cast (WPI) and was then fixed on skull with dental adhesive resin cement (Super-Bond, SUN MEDICAL).

### *Ex vivo* electrophysiology

Mice were deeply anesthetized with isoflurane and decapitated. Horizontal sections (300 µm) containing ventral hippocampus were prepared by vibratome (VT1200S, Leica) in ice-cold choline solutions containing (in mM): 25 NaHCO_3_, 1.25 NaH_2_PO4-H_2_O, 2.5 KCl, 0.5 CaCl_2_, 7 MgCl_2_, 25 D-glucose, 110 Choline chloride, 11.6 Ascorbic acid, and 3.1 C_3_H_3_NaO_3_. Slices were subsequently incubated in artificial cerebrospinal fluid (ACSF) containing (in mM): 125 NaCl, 25 NaHCO_3_, 1.25 NaH_2_PO4-H2O, 2.5 KCl, 11 D-glucose, 2 CaCl_2_, and 1 MgCl at 34 °C for 30 mins then at room temperature (∼20 °C) for at least 1 hour before the recording.

Whole-cell recording was performed with an upright microscope (BX51WI, Olympus) equipped with a motor-controlled stage and focus (MP285A and MPC-200, Sutter instrument), differential interference contrast, coolLED (model pE300), and a monochrome camera (Moment, Teledyne). Neurons were visualized with a 60x lens (1.00 NA, LUMPlanFL N, Olympus) with the software Micro-Manager-2.0 gamma (US National Institutes of Health). Pipettes (4-5 MΩ) were pulled from thick-walled borosilicate capillary glass with a puller (model P-1000, Sutter instrument). For a voltage-clamp recording, the internal solution contained (in mM): 135 CsMeSO_3_, 10 HEPES, 4 Mg-ATP, 0.3 Na-GTP, 8 Na_2_-Phosphocreatine, 3.3 QX-314 (pH was adjusted to 7.35 with CsOH). In some experiments, a biocytin (4 mg/mL) and Alexa488/586 (50 μM) were further added for morphological studies. Recordings and hardware control were performed with Wavesurfer (Janelia Farm). Signals were amplified and Bessel filtered at 4 kHz with an amplifier (MultiClamp 700B, Molecular Device), then sampled at 10 kHz with a data acquisition board (USB-6343, National Instrument). Recording with access resistance change of > 20% from the baseline (∼30 MΩ or less) were discarded. Liquid junction potential was not corrected. All recordings were performed at ∼34 °C maintained with the inline heating system (TC-324C, Warner instrument).

To stimulated channelrhodopsin-2, a blue LED was delivered at a short pulse (5-msec) through a 4x objective lens (0.16 NA, UPlanSApo, Olympus). The light power (measured at the focal plane) was 20 mW for the recordings in normal ACSF and 40 mW for the recordings in the presence of tetrodotoxin and 4-aminopyridine. Photo-stimulation was repeated 3 to 6 times (at intervals of 10 or 20 sec) to obtain an average response trace. Photo-evoked responses less than 2 standard deviations of the baseline were considered as a “zero” response. The mean amplitudes indicate mean amplitudes between LED stimulus-onset to 50 ms of average responses.

Drugs used for ex vivo electrophysiology were purchased from Merck, TOCRIS, and Hellobio.

### Quantification of presynaptic neurons

Mice were deeply anaesthetized with a ketamine (120 mg/kg) and xylazine (24 mg/kg) mixture and were transcardially perfused with 4% paraformaldehyde (PFA). The brains were extracted and immersed in 4% PFA overnight followed by 20 and 30% sucrose solution for a cryoprotection. The brains were subsequently embedded in O.C.T compound (Tissue TEK, Sakura) in a mold and frozen on the dry ice. Coronal slices (100 µm) of entire brain were cut with Cryostat (CM3050S, Leica) and were washed with phosphate-buffered saline (PBS) before DAPI staining. All sections were mounted onto glass slides with a coverslip (thickness:0.13-0.17 mm, Hounisen) and mounting media (DAKO, S3023, Agilent Technologies).

The injection sites and its specificity were verified by registering the images into Allen Brain atlas using QuickNII and VisuAlign (*57*).

For the visualization and quantification of presynaptic neuron distribution in a 3D mouse brain, epifluorescence images were acquired using a Slidescanner (4231 x 3462 pixels, 10x objective, Olympus VS120) and analyzed with the MATLAB-based program AMaSiNe (*34*). Specifically, for **Fig. 1**, coronal sections between AP +3 mm and −4 mm were registered in AMaSiNe, which semi-automatically detects fluorescently labeled presynaptic cells across all sections. To control for transfection variability (which may arise from slight difference in injection volume or site), we normalized the number of presynaptic neurons in the specified brain areas (shown in **Fig. 1H**) by the total number of presynaptic neurons in those regions.

For the monosynaptic rabies tracing study in **Fig. 6**, the brain areas containing over 1% presynaptic cells of total presynaptic cells were proceeded further analysis. Fraction of presynaptic cells in the different areas were calculated by dividing a number of presynaptic cells in each brain areas by the average number of presynaptic neurons in all brain areas.

For the study in **Fig. 2**, the sections were immunostained with NeuN before imaging. Here, sections were immersed with 5% normal goat serum and 0.2% Triton X for 30 mins at room temperature before incubation with NeuN primary antibody (anti-rabbit, 1:1000) (ABN78, Milipore) at 4 °C for overnight. The sections were then washed with PBS and were further incubated with secondary antibody (goat anti-rabbit conjugated with Alexa 488 or 568, 1:500) (A11008 or A11036, Thermo Fisher) for 1 hour at room temperature before mounting. A single plane image of CLA complex was acquired with confocal microscope (10x objective, 1024 x 1024 pixels, Zeiss). 2-4 images, range of AP: +1.4 to −0.2, were collected per brain. Fraction of double-labelled neurons (with different retrograde tracers) were calculated for each brain by dividing a total number of double-labeled neurons by the total number of vCA1-projecting neurons (labeled with one color).

### Quantification of fluorescence signal from axons

For **Fig. 3 and S3**, horizontal section (100 µm) images were taken with Sliderscanner (10x lens, pixel: 2560 x 3072). All images from the same brain were taken with the same light intensity and detection setting. After background subtraction, GFP signals in ventral, intermediate, and dorsal CA1 column were normalized by the highest signal in ventral CA1 in the same brain.

For **Fig. 2**, coronal sections (100 µm) were immunostained with GFP antibody before imaging. Here, sections were washed with PBS, the sections were incubated in blocking solution containing 5% normal goat serum and 0.2% Triton X for 30 mins at room temperature before incubation with GFP primary antibody (anti-rabbit, 1:1000) (NB600-308, MNovus Biologicals) in blocking solution at 4 °C for overnight. The sections were then washed with PBS and were further incubated with secondary antibody (goat anti-rabbit conjugated with Alexa 488, 1:300) (A11008, Thermo Fisher) for 1 hour at room temperature before mounting. Images were taken with Slidescanner (10x, pixel: 3072 x 3584). Signal per pixel in ROI was calculated for each projection site after background subtraction.

### Social memory test with chemogenetic inhibition of EN^vCA1-proj.^ neurons

Social memory test is based on previous articles with slight modifications (*58*), (*59*). All tested mice were habituated to the handling for 3 days. The water-soluble CNO purchased from Hellobio was dissolved by saline (final concentration was 0.1mg/ml). Prior to the test day, we divided subject mice into two groups. On the 1^st^ test day (session 1), one group received chlozapine-N-oxide (CNO, 1 mg/kg, i.p.) and another group received a vehicle. After 30 mins, the subject mice were placed in a three-chamber box for 5 mins (habituation), then the same box with two pencil chambers containing the same object in opposite corners for 5 mins (pretest). After 5 mins break in their home cage, the subject mice were again placed in the same box, but this time, an object in one pencil chamber was replaced with different object and an object in another pencil chamber was replaced with a conspecific (3-4 weeks old male, habituated to the handling and pencil chamber for 3 days prior to the session 1). This “sociability test” lasted for 10 mins and was followed by social discrimination test after 30 mins break in the home cage. The social discrimination test lasted for 5 mins and consisted of the same setting as the sociability test except an object in a pencil chamber, which was replaced with a new conspecific (i.e., a novel mice). In the following day, CNO and saline treatments were swapped between the groups, and all mice went through the same series of tests (session 2) for the within-subject comparison of the CNO effect. The position of the pencil chamber including an object or a conspecific were counterbalanced. The tests were performed under 6-10 lx room light and video was recorded at 25 frames/sec with GigE camera (Basler ace acA1300-60gm, Basler) placed at the ceiling.

### Trace fear conditioning

After one week of social memory test, subject mice were examined for their tone-associated fear memory and extinction. One group received CNO while another group received vehicle 45 mins before the test. In conditioning day, mice were placed in a box with a metal grid floor and ethanol odor (context A) and after 3 mins were delivered with 20-sec pure tone (2 kHz, 20 dB) followed by 18-sec time gap (trace) and 2-sec footshock (0.5 mA). The exposure to the same tone-trace-shock sequence was repeated 2 more times with inter-trial interval of 2 mins. In the following day, mice were re-exposed to the context A for 3 mins to assess their fear memory to the context. From day 3, mice were placed in a new context, the context B (smooth floor, breach odor), and went through the same protocol as in conditioning day but without shocks. The process was repeated over 5 days to assess the extinction of the fear memory to the tone and trace. All test was performed in dark (but infrared light was on), and behavior was video recorded at 25 frames/sec with infrared GigE camera (Basler ace acA1300-60gm, Basler).

### Novel object recognition test

The novel object recognition test is based on the previous article with slight modifications (*58*). The subject mice were divided into two groups and CNO treatment was done as same as social memory test. 40 mins after CNO/vehicle i.p. injection, the subject mice were placed in an open field box (50 x 50 cm) for 5 mins (habituation). In following familiarization session, two same shape objects are placed in the open field box, and then the subject mice explored for 10 mins. After 1 hour break in their home cage, the subject mice were again placed in the same box, but one object is replaced by a novel object. The subject mice are allowed to explore the box for 5 min in discrimination session.

### Social memory test and novel object recognition test with *in vivo* fiber photometry

Optic probe implanted mice were habituated handling for 3 days prior the test. In test day, the subject mouse attached with patch code were placed in an open field box (50 x 50 cm) for 5 min (habituation). After habituation, the mouse was place in a home cage for resting and waiting following session. For pretest, pencil chambers containing the same objects were placed at the corners of the open field box. Pretest was tested for 5 min. For sociability test, a novel mouse (4 weeks old, male) or a novel object were placed in pencil chambers. Sociability test took 10 min. After sociability test, the subject mouse return to a home cage and rest for 20 min. After 20 min rest, the subject mouse was tested discrimination session for 5 min. In discrimination test, a novel mouse (4 weeks old, male) or a familiar mouse that was used as the stimulus mouse in sociability test were placed in pencil chambers. Both in sociability and discrimination test, positions of pencil chambers were counterbalanced between subject mice. The tests were performed under 6-10 lux room light and video was recorded at 60 frames/sec with GigE camera (Basler ace acA1300-60gm, Basler) placed at the ceiling. Video acquisitions were controlled by Bonsai. Simultaneous start of video recording and calcium signal recording was controlled by Raspberry Pi.

After one week of social memory test, subject mice were examined for novel object recognition test. The procedure, the arenas and objects were used the same as in the novel object recognition test with chemogenetic inhibition of EN^vCA1-proj.^ neurons.

### Behavior data acquisition and analysis

Ethovision (version 16 or 17, Noldus) was used for the hardware control and data acquisition and initial quantification for the behavior. Behavioral variables, including movement trajectory, times spent in predefined areas, total distance traveled, travelling velocity, and freezing were extracted and further analyzed/nested with custom functions in the Matlab (R2020b). For social memory test, interaction zones were defined as an area between the edge of pencil chamber and 3 cm away from the edge. When the nose of the subject mouse was in the zone, the time was counted as interaction time. Sociability index was calculated by dividing total interaction time with mouse by total interaction time with mouse and object. The discrimination index was calculated by dividing total interaction time with a novel mouse by total interaction time with a familiar mouse and a novel mouse. Data from mice were excluded if discrimination index was less than 0.5 during saline treatment. For trace fear conditioning test, mice were considered “freezing” if pixel change between frames was less than 0.1 % for more than 1 sec.

Deeplabcut (ver.2.3) was used for tracking of annotated body points of the subject mice connected with patch code for calcium imaging of EN^vCA1-proj.^ neurons during social memory test and novel object recognition test. Nose points were used for further analysis using Matlab (R2020b). To define interaction zone, reference points was set at the center of pencil chamber.

The areas that distance between nose point and reference point was less than 100 pixel for social memory test and 50 pixel for novel object recognition test were dedicated to interaction zone.

### Fiber photometry and analysis

Population activity of EN^vCA1-proj.^ neurons were recorded using fiber photometry (Doric Lenses). GCaMP8s was excited with 470 nm light at 20-30 μW measured at the tip (isosbestic point used was 415 nm (15 μW)). Calcium-dependent and independent signals were collected using lock-in mode. Signals were detected at 10x gain. Recorded signals were butterworth filtered (low-pass at 40Hz) and calcium signals (in z-score) were extracted using Doric neuroscience studio V6 (Doric Lenses). Signals were further down sampled to 60 Hz using Matlab (R2020b) signal analyzer to match the behavioral sampling data (60 fps). Z-score > 2.56 (alpha = 0.01) was determined to be a calcium event. For heat map of cumulative time and calcium events, spatial binning was 50 pixels. To make correlation map (**Fig. S7 and S8A**), correlations between column of cumulative time heat map and cumulative calcium event heat map were tested using Matlab corr function.

### Statistical analysis

Data and statistical analysis were performed with MATLAB R2020b (Mathworks). Unless otherwise noted, group data was presented as median, and first and third quartiles are shown for dispersion. Statistics were performed using a non-parametric test (signed-rank test for paired data and rank-sum test for unpaired data). One-way ANOVA (post hoc test: Tukey-Kramer test or Kruskal-Wallis test) were applied for multi-group comparison. Two-way ANOVA were performed for comparison of multi-factors. Pearson correlation coefficient was applied for correlation analysis. Significance was defined as *p < 0.05, **p < 0.01, ***p < 0.001. Outlier was detected by Grubbe test (GraphPad, significant level: 0.05). Sample number were indicated as “n”, animal number were indicated as “N”.

## Acknowledgments

We thank Wen-Hsien Hou, Chihiro Nakamoto, Ana Cicvaric, Hui Zhang, and Elizabeth Wood for constructive discussion of this study. We thank Peter Bjerge, Bjark B. Brix, and Dennis Olesen for technical help. We acknowledge the bioimaging core facility, Health, Aarhus University, Denmark, for the use of equipment and support.

## Funding

LF professorships, R310-2018-3611 (JR) LF experiment, R436-2023-471 (NY) LF experiment, R436-2023-855 (AT)

## Author contributions

Conceptualization: AT

Methodology: AT, NY, BMM

Investigation: AT, NY, HL, SJF, BMM, MZ, MM

Supervision: AT, NY, JR

Writing—original draft: AT, NY

Writing—review & editing: AT, NY, JR

## Competing interests

The authors declare no competing interests.

## Data and materials availability

All data needed to evaluate the conclusions in the paper are present in the paper and/or Supplementary Materials.

## Supplemental figures

**Fig. S1.**
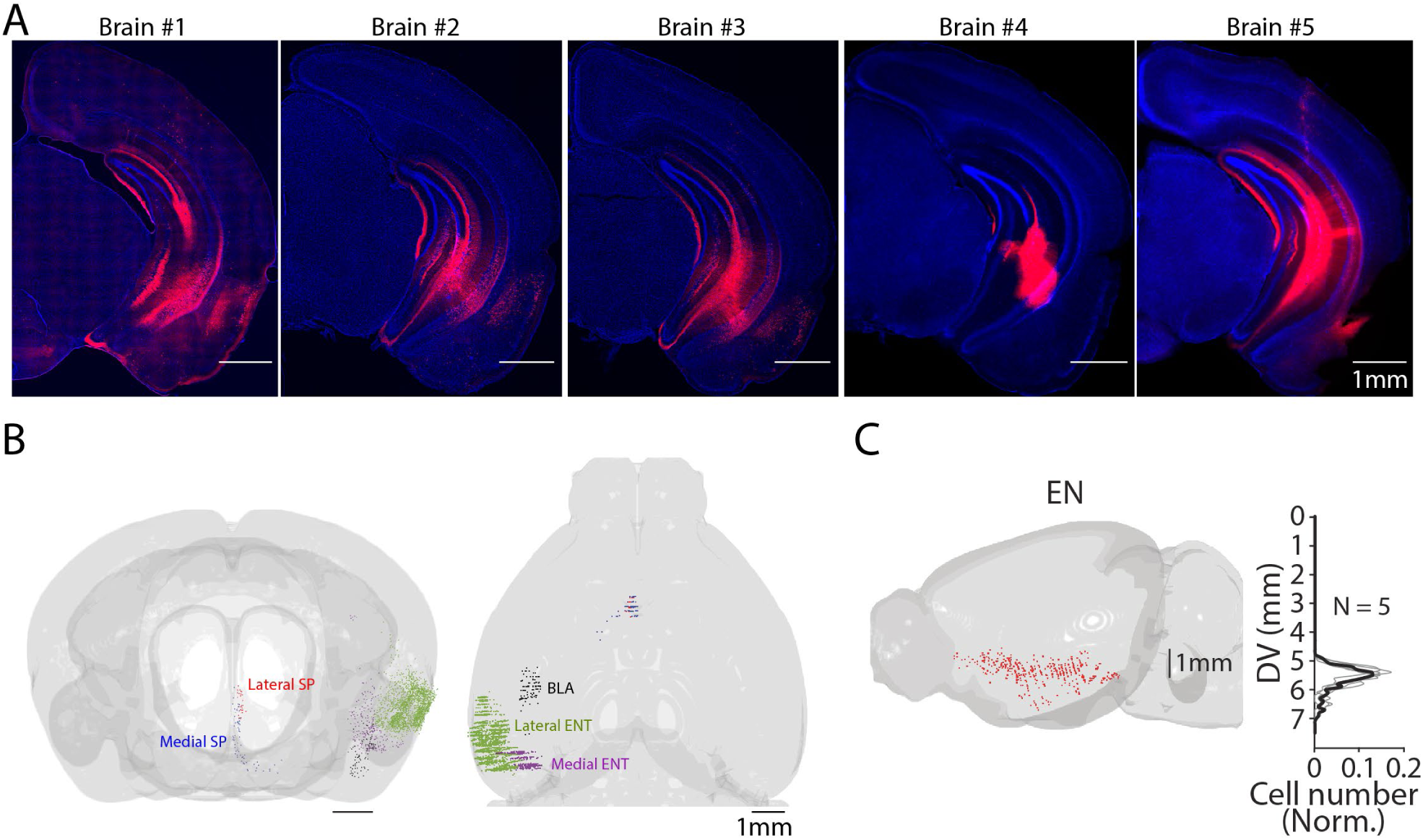
Whole brain distribution of vCA1-projecting neurons. (**A**) Fluorescent images of the injection site (vCA1) of CAV2-Cre (60 nl) of Ai14 mice for analysis of presynaptic neuron distribution using AMaSiNe. The presynaptic cell numbers counted in individual brain are presented in Table 1. (**B**) An example distribution map of vCA1-projecting neurons in SP (lateral and medial), BLA and ENT (lateral and medial) shown from front and top view. Neurons in different brain areas are color-coded. (**C**) The side view of the brain shown in Fig. 1J. Plots indicate distribution of vCA1-projecting neurons in EN along dorsal-ventral axis (DV). Data from different mice are shown in gray. Median is indicated in black.

**Fig. S2.**
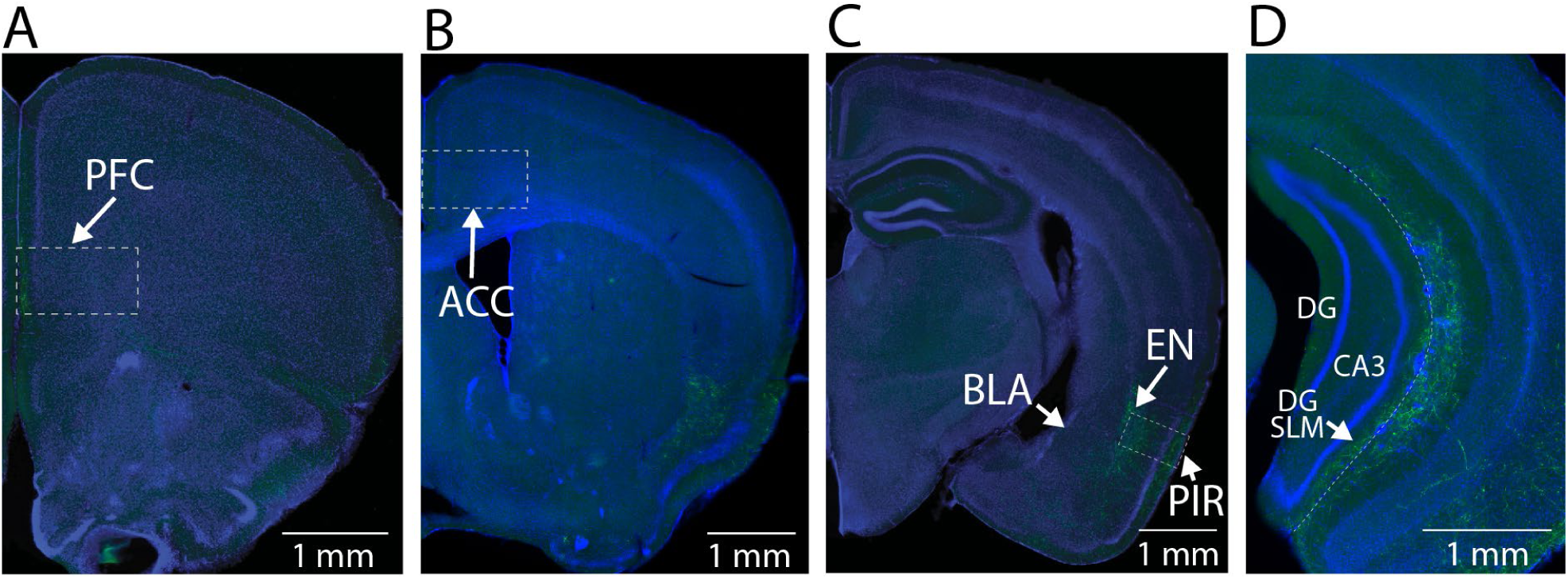
Axon branching of vCA1-projecting EN neurons. (**A-D**) Example fluorescent images of branching of vCA1-projecting neurons. All images were from the same mouse shown in Fig. 2N-O. DG: Dentate gyrus

**Fig. S3.**
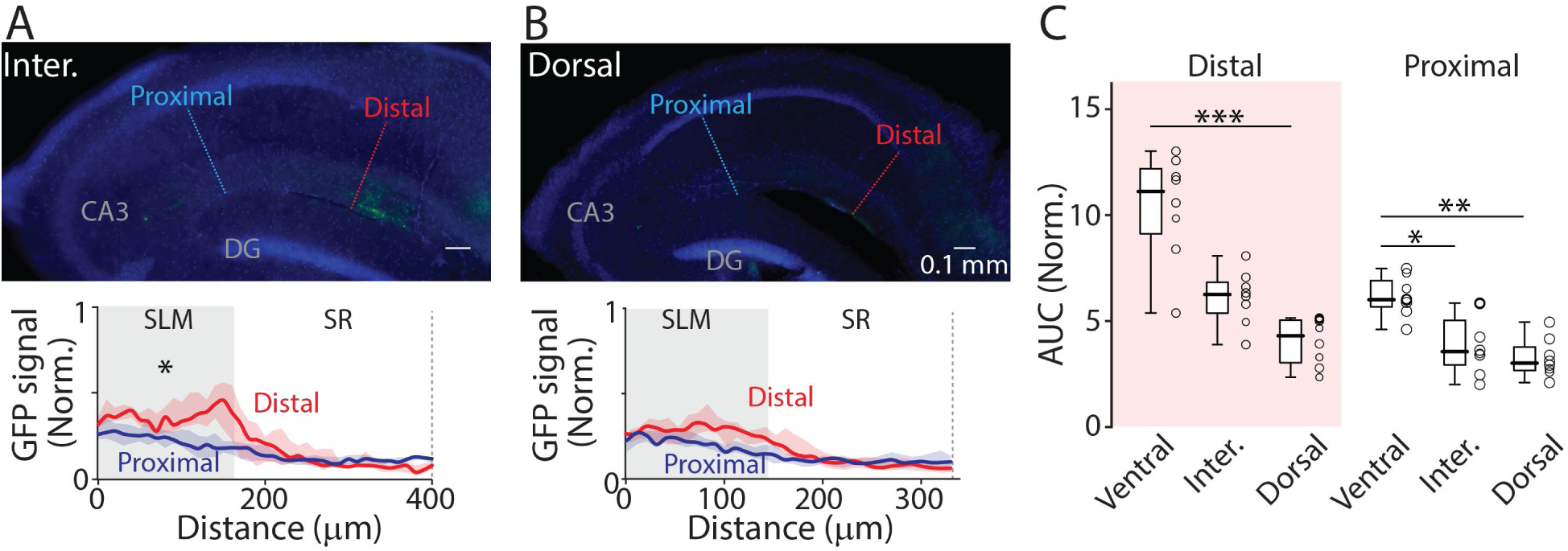
Projections of EN^vCA-proj.^ neurons to the hippocampus. (**A**) Top: An example fluorescent images of intermediate hippocampal sections from the same brain as in Fig. 3B and C. Bottom: GFP signal profile at the distal and proximal region indicated in the top image. (**B**) The same as in (**A**) but for dorsal hippocampus. Median area under the curve (AUC): distal vs proximal (intermediate), 6.3 vs 3.5, p = 0.007; distal vs proximal (dorsal), 4.3 vs 3.0, p = 0.055; Wilcoxon signed-rank test. n = 8 slices (N = 4 mice). (**C**) Comparison of GFP signal (in AUC) between different hippocampal axis for distal or proximal vCA1. Distal; ventral vs intermediate, p = 0.086; ventral vs dorsal, p < 0.001; intermediate vs dorsal, p = 0.136. Proximal: ventral vs intermediate, p = 0.018; ventral vs dorsal, p = 0.001; intermediate vs dorsal, p = 0.695; Kruskal-Wallis test. n = 8 slices (N = 4 mice).

**Fig. S4.**
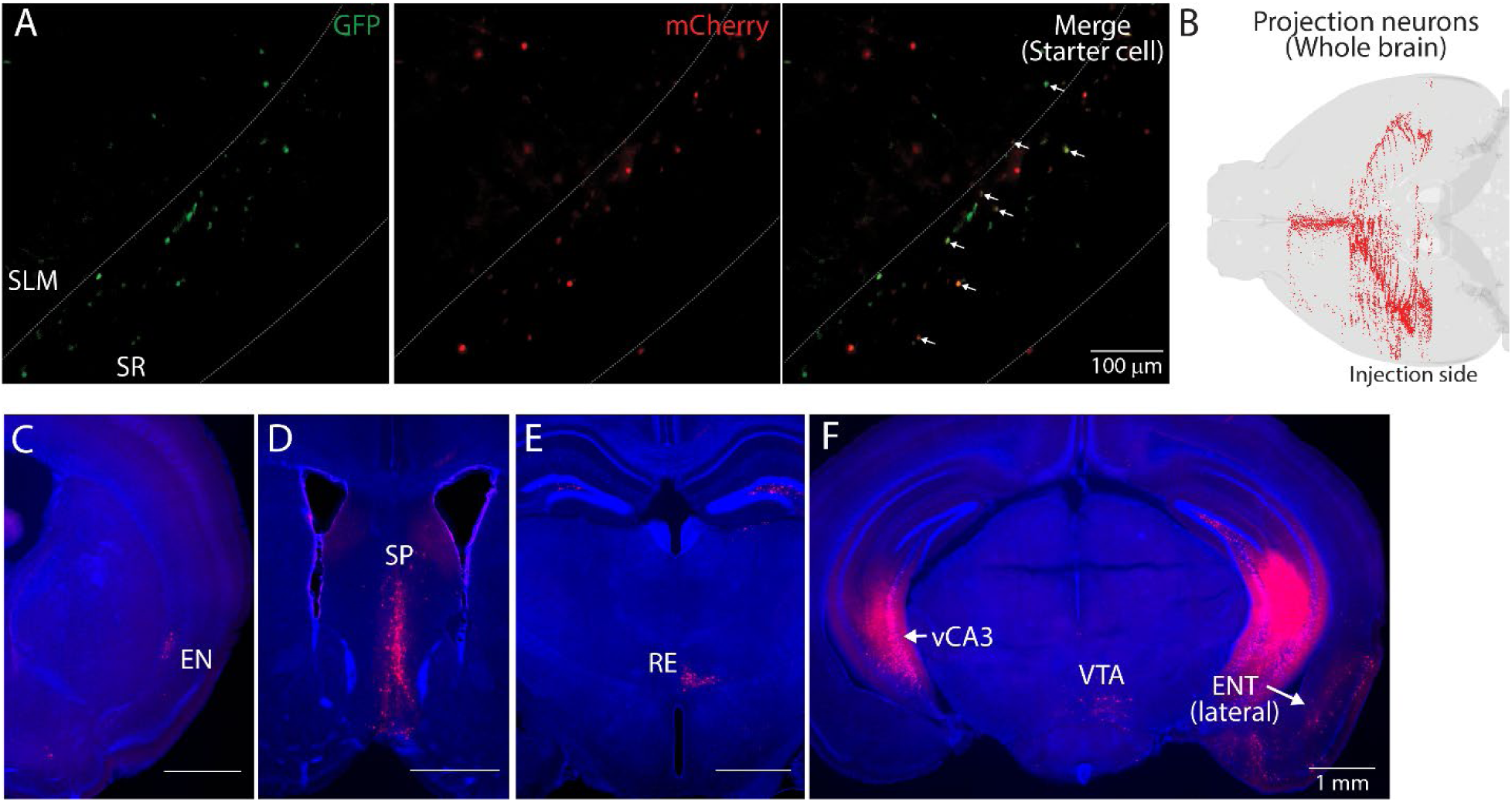
Presynaptic neurons of ventral hippocampal GABAergic neurons. (**A**) Example fluorescent images of starter cells in vCA1. TVA_oG expressing neurons were labeled by GFP and rabies expressing neurons were labeled by mCherry. GFP and mCherry double expressing neurons were counted as starter cells. White allows in eight panel indicate starter cells. White dashed lines indicate the Py-SR and SR-SLM border. (**B**) An example map of presynaptic cells in a whole brain. Injection was made in the left hemisphere. (**C-F**) Fluorescent images of presynaptic cells in coronal sections from the same mouse brain. Presynaptic neurons were in EN (**C**), septum(SP) (**D**), nucleus reunion (RE) (**E**), contralateral vCA3 (vCA3) (**F**), ventral tegmental area (VTA) (**F**), and lateral ENT (**F**).

**Fig. S5.**
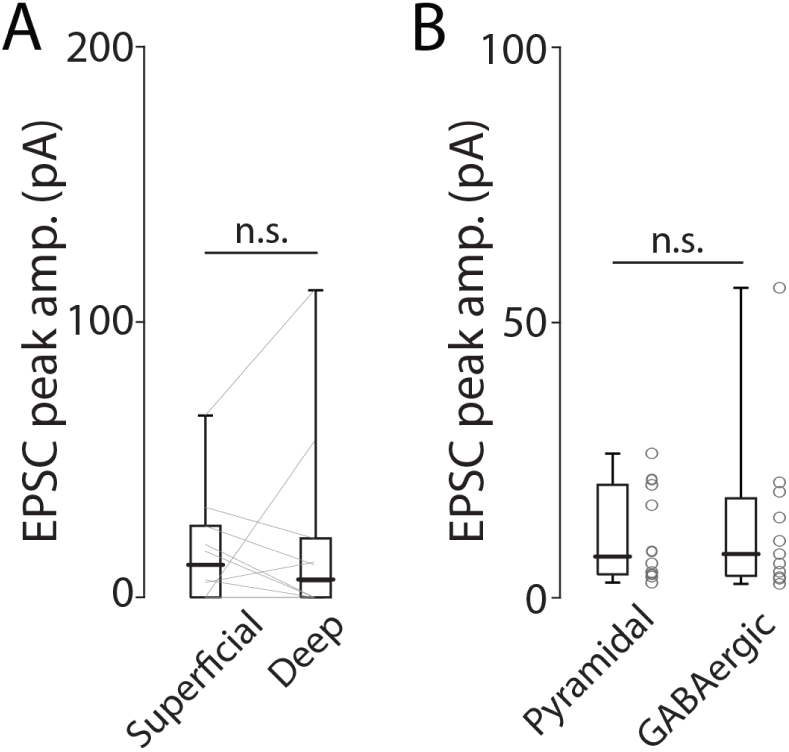
EN-mediated excitatory inputs to ventral hippocampal neurons. (**A**) Plots comparing peak amplitude of photo-evoked EPSCs in superficial and depp pyramidal neurons. Thin gray lines indicate paired data. Median peak amplitude; superficial vs deep (in pA): 11.5 vs 5.9, p = 0.625; signed-rank test. (**B**) Plots comparing peak amplitude of photo-evoked EPSCs in pyramidal and GABAergic neurons. The data from superficial and deep pyramidal neurons were pooled. n = 10 for pyramidal neurons and 11 for GABAergic neurons. N = 4 mice. Median EPSC peak amplitude: Pyramidal vs GABAergic (in pA): 7.4 vs 8.0, p = 0.86. Rank-sum test.

**Fig. S6.**
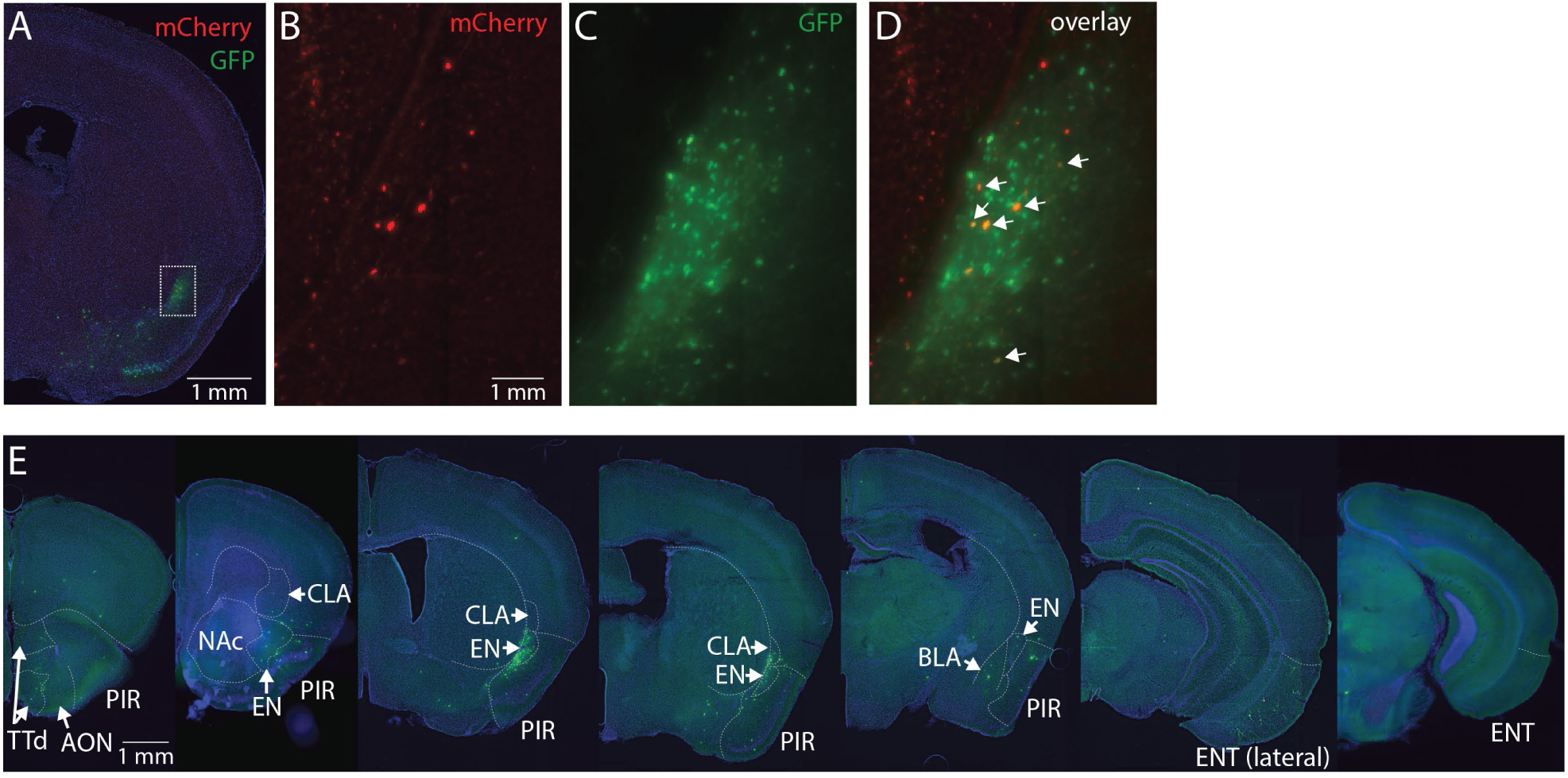
Presynaptic neurons of EN^vCA1-proj.^ neurons. (**A**) An example fluorescent image of starter cells in EN. (**B-D**) Expanded images of the area marked by white square in panel A. White allows indicate starter cells. (**E**) Example fluorescent images of coronal sections in series. Labeled neurons (in green) indicate presynaptic cells of EN^vCA1-proj.^ neurons. TTd: Taenia tecta, AON: Anterior olfactory nucleus, PIR: Piriform cortex, NAc: Nucleus accumbens, EN: Endopiriform, CLA: Claustrum, BLA: Basolateral amygdala, ENT: Entorhinal cortex.

**Fig. S7.**
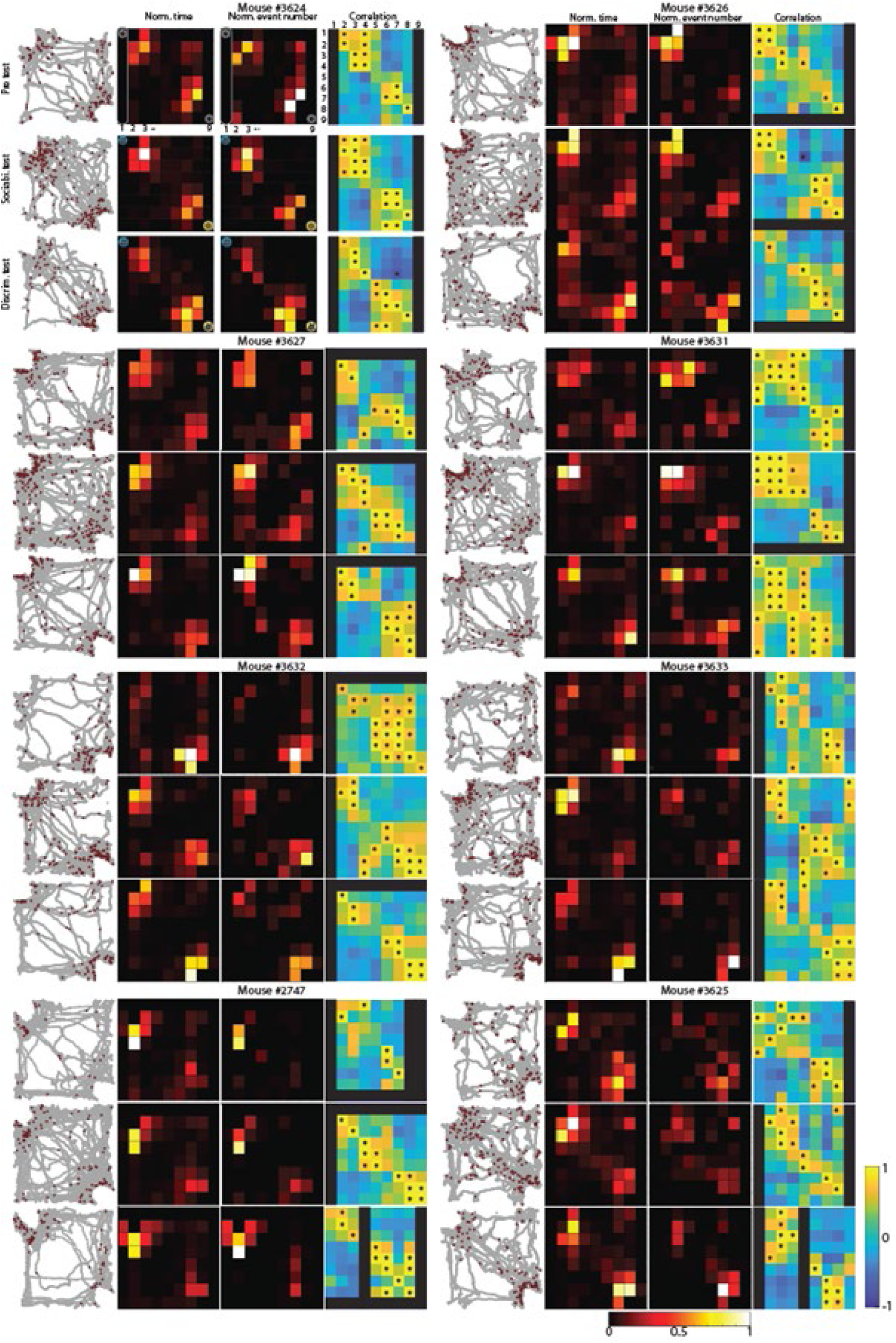
Nose point location and calcium signal data during social memory test. Maps of nose points (filled gray circles) and calcium events (filled red circles) in arena. Heat maps of the same data (but cumulative time and calcium event are separated) were also shown for each mouse (N = 8). Peason’s correlation coefficients with p < 0.05 is indicated by asterisk. All p values were presented in table S1.

**Fig. S8.**
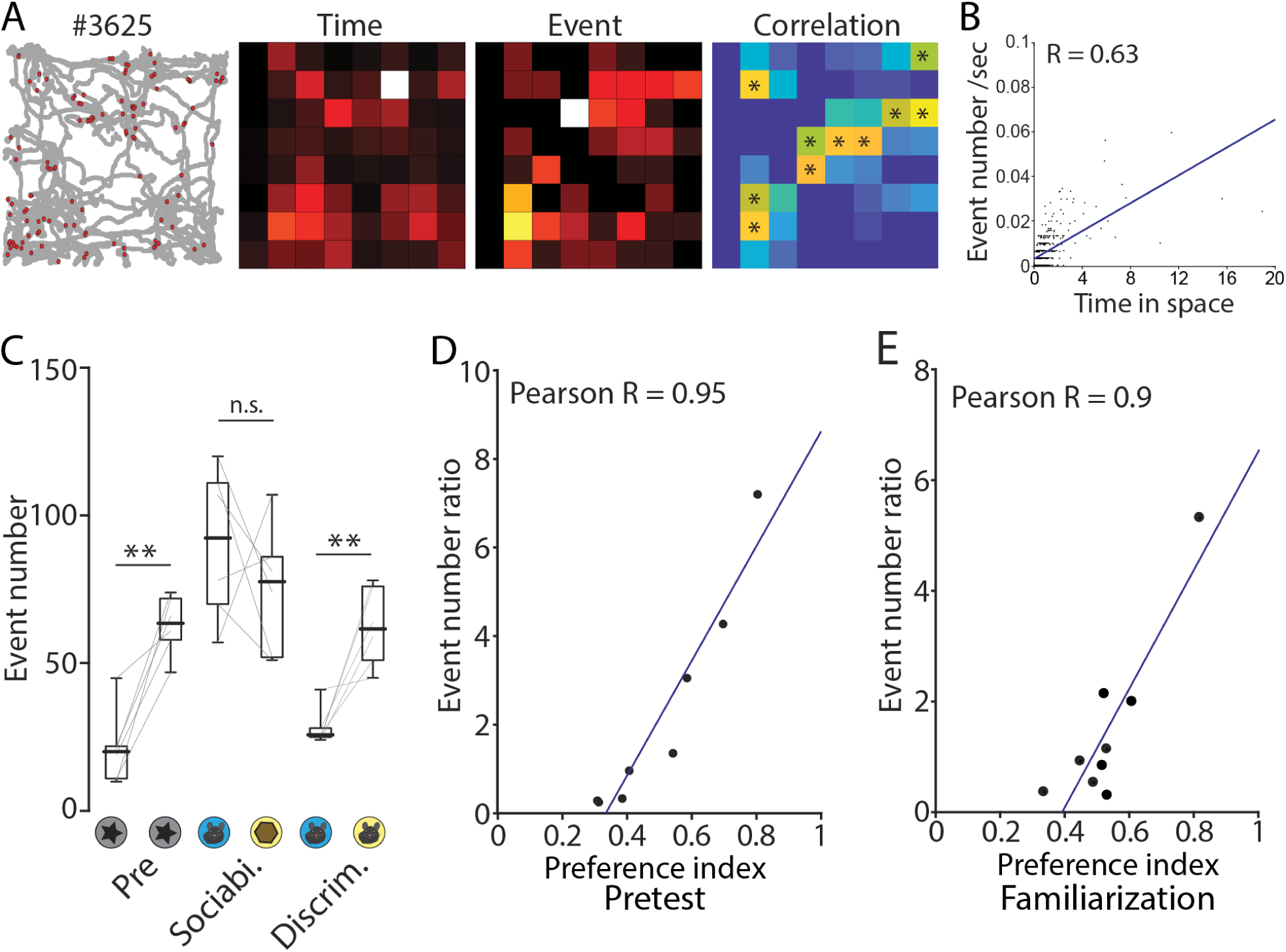
Calcium signals during exploratory behavior in open field, pretest and familiarization test. (**A**) Example of maps of nose points and calcium events in open field arena. All p values in a correlation map were presented in table S2. (**B**) Correlation plots of event number and cumulative time in each pixels shown in A. p < 0.001, N = 7. (**C**) Box plots comparing a calcium event number when the mouse was in the interaction zones. n the pretest plot, the object that the mice interacted with more is placed on the right side. Median event number: Pre (left vs right), 20 vs 63.5; Sociability (mouse vs object), 92.5 vs 77.5; Discrimination (familiar vs novel), 22.5 vs 63.5. p = 0.001, 0.385, and 0.006. Signed-rank test. N = 6. (**D** and **E**) Correlation plots of calcium event number ratios and preference index in pretest (**D**) and familiarization session (**E**). Pretest, p < 0.001, N = 8; Familiarization, p = 0.001, N = 9.

**Fig. S9.**
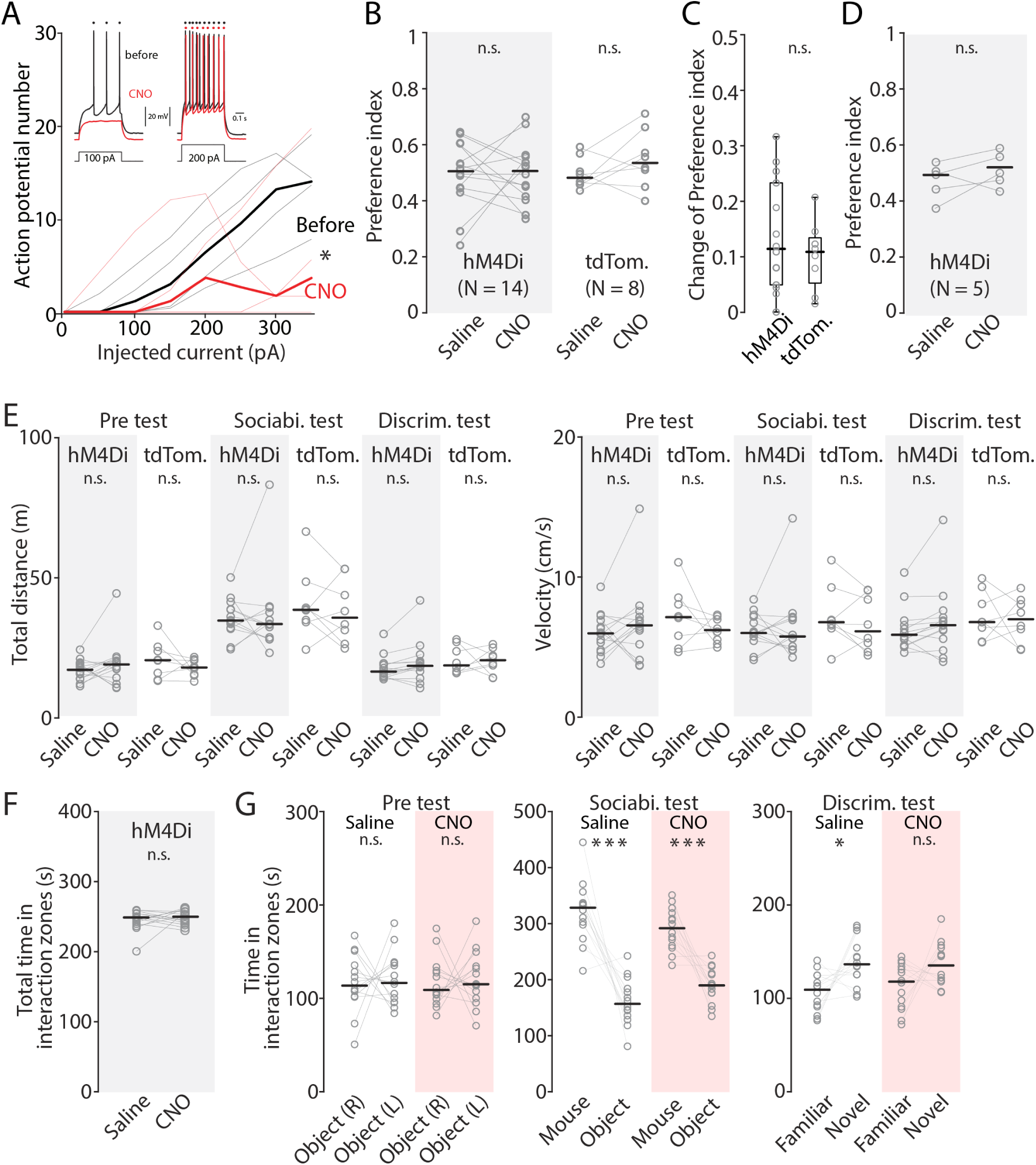
The effect of chemoegentic inhibition of EN^vCA1-proj.^ neurons on the pretest and locomotion. (**A**) Frequency-current (FI) curve showing the effect of CNO on action potential number generated by current injection. Recording was made from hM4Di expressing EN^vCA1-proj.^ neurons before and after bath application of CNO (100 nM). Black line indicates before and red line indicates after CNO application. Inset: Traces showing spike response to 100 pA (left) and 200 pA (right) current injection. Saline vs CNO; F (1, 4.77) = 0.38, p = 0.0329, two-way ANOVA. n = 4 neurons. N = 2 mice. (**B**) Same as in Fig. 8E, but for preference index in pretest. Median preference index: hM4Di mice (Saline vs CNO), 0.5 vs 0.5, p = 0.92; signed-rank test; tdTomato mice (Saline vs CNO), 0.48 vs 0.53, p = 0.227; signed-rank test. (**C**) Absolute change of preference index before and after CNO treatment in hM4Di and tdTomato mice. Median change of preference index: hM4Di mice 0.11 (N = 14); tdTomato 0.11 (N = 8). hM4Di vs tdTomato, p = 0.517, ranksum test. (**D**) Same as in B, but for preference index in familiarization session of novel object recognition test. Median preference index: hM4Di mice (Saline vs CNO), 0.49 vs 0.5, p = 0.28; signed-rank test (**E**) Plots comparing locomotor activity (distance moved and velocity) for hM4Di and tdTomato mice under two different treatments. Median total distance: Pretest, hM4Di mice (Saline vs CNO), 17.17 vs 19.10, p = 0.426; tdTomato mice (Saline vs CNO), 20.60 vs 17.98, p = 0.25; Sociability test, hM4Di mice (Saline vs CNO), 34.72 vs 33.51, p = 0.855; tdTomato mice (Saline vs CNO), 38.59 vs 35.76, p = 0.547, Discrimination test, hM4Di mice (Saline vs CNO), 16.55 vs 18.58, p = 0.0676; tdTomato mice (Saline vs CNO), 18.73 vs 20.60, p = 0.945, all signed-rank test. Median velocity: Pretest, hM4Di mice (Saline vs CNO), 5.98 vs 6.55, p = 0.391; tdTomato mice (Saline vs CNO), 7.14 vs 6.21, p = 0.383; Sociability test, hM4Di mice (Saline vs CNO), 6.01 vs 5.76, p = 0.808; tdTomato mice (Saline vs CMO), 6.78 vs 6.13, p = 0.313; Discrimination test, hM4Di mice (Saline vs CNO), 5.88 vs 6.57, p = 0.104; tdTomato mice (Saline vs CNO), 6.79 vs 6.99, p = 0.945. All signed-rank test. (**F**) Total time in interaction zones during discrimination test (familiar mouse interaction zone + novel mouse interaction zone). Median total time: (Saline vs CNO), 248.32 vs 294.37; p = 0.67, signed-rank test. (**G**) Time in interaction zones during pre test, sociability test, and discrimination test in hM4Di mice with saline or CNO treatment. Median time in interaction zone: Pretest, Saline (object (in right chamber) vs object (in left chamber)), 116.25 vs 120.26, p = 1, CNO (object (in right chamber) vs object (in left chamber)), 117.29 vs 120.2, p = 0.855; Sociability test, Saline (Mouse vs Object), 320.89 vs 161.2, p < 0.001, CNO (Mouse vs Object), 290.59 vs 189.75, p < 0.001; Discrimination test, Saline (Familiar vs Novel), 106.35 vs 138.71, p = 0.02, CNO (Familiar vs Novel), 113.1 vs 135.69, p = 0.1353. All signed-rank test.

**Fig. S10.**
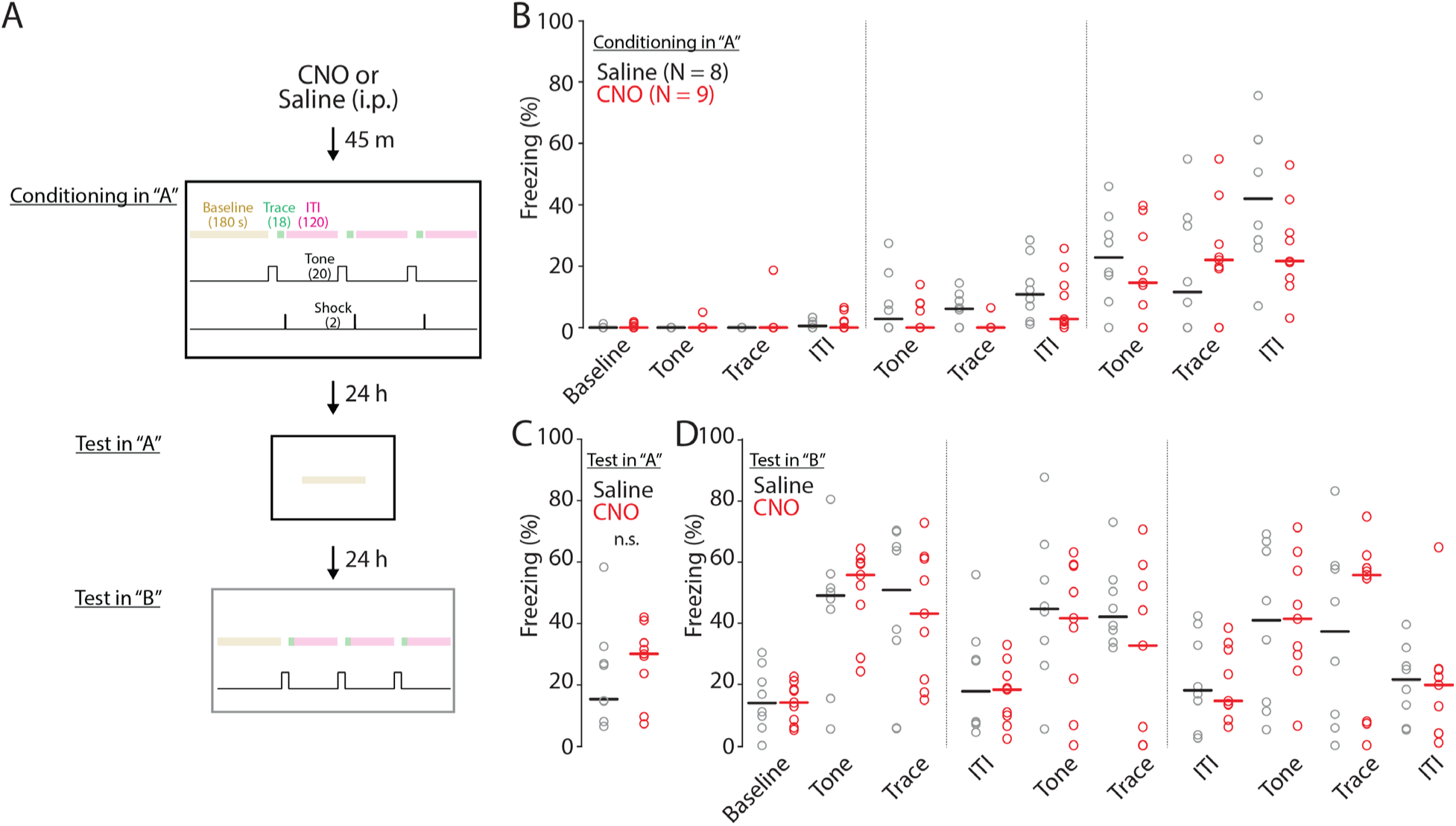
The effect of chemoegentic inhibition of EN^vCA1-proj.^ neurons on trace fear conditioning. (**A**) Schematic of trace fear conditioning protocol. (**B**) The effect of CNO on freezing behavior during conditioning. Black indicates saline treated group and red indicates CNO treated group. Data points of individual mice are indicated by empty circles. Thick horizontal bars indicate median values. Saline vs CNO, F(1, 9) = 2.52, p = 0.115; two-way ANOVA. (**C**) The effect of CNO on freezing behavior during recall to context “A”. Median values of freezing %: Saline = 14.91, CNO = 30.13. Saline vs CNO, p = 0.576; one-way ANOVA. (**D**) The effect of CNO on freezing behavior during conditional stimulus recall in context “B”. Saline vs CNO, F (1, 9) = 0.38, p = 0.54; two-way ANOVA

## Notes

### Competing Interest Statement

The authors have declared no competing interest.

### Summary of Updates

New data set related behavior tests were added as Fig. S9G. The text was edited to clarify the findings.

